# The dynamics of capillary flow in an open-channel system featuring trigger valves

**DOI:** 10.1101/2024.09.17.613325

**Authors:** Jodie C. Tokihiro, Ingrid H. Robertson, Denise Gregucci, Albert Shin, Elisa Michelini, Tristan M. Nicholson, Ayokunle Olanrewaju, Ashleigh B. Theberge, Jean Berthier, Erwin Berthier

## Abstract

Trigger valves are fundamental features in capillary-driven microfluidic systems that stop fluid at an abrupt geometric expansion and release fluid when there is flow in an orthogonal channel connected to the valve. The concept was originally demonstrated in closed-channel capillary circuits. We show here that trigger valves can be successfully implemented in open channels. We also show that a series of open-channel trigger valves can be placed alongside or opposite a main channel resulting in a layered capillary flow. We developed a closed form model for the dynamics of the flow at trigger valves based on the concept of average friction length and successfully validated the model against experiments. For the main channel, we discuss layered flow behavior in the light of the Taylor-Aris dispersion theory and in the channel turns by considering Dean theory of mixing. This work has potential applications in autonomous microfluidics systems for biosensing, at-home or point-of-care sample preparation devices, hydrogel patterning for 3D cell culture and organ-on-a-chip models.

## Introduction

Microfluidic devices precisely move fluids through small channels and can use surface tension effects (capillary forces) defined by channel geometry and surface chemistry to achieve self-powered and self-regulated operation. Capillary microfluidics is driven through spontaneous capillary flow (SCF)^1–7^ and can perform timed multi-step processes by leveraging capillary forces encoded in device architecture without the need for external triggers (e.g., pushing a button, programming an electrical signal, or other user activity).^8–11^ Trigger valves (TGVs) are one of the main geometric features/control elements that make autonomous capillary-driven possible. TGVs are modified passive stop valves that release a confined liquid upon the capillary-driven flow of another or a similar liquid in an orthogonal channel to the stop valve (Figure 1A). These valves are widely used in a variety of closed-channel point-of-care diagnostic applications such as immunoassays for bacterial, antibody, and protein detection antibody or protein detection as well as live-cell staining.^12–15^ There is a wealth of theoretical, experimental, and applied work using closed-channel TGVs.^12–24^ While the concept of using TGV in open microfluidic systems have been introduced in brief, ^19,25,26^ there is a need for more in-depth theoretical development and experimental validation.

**Figure 1.**
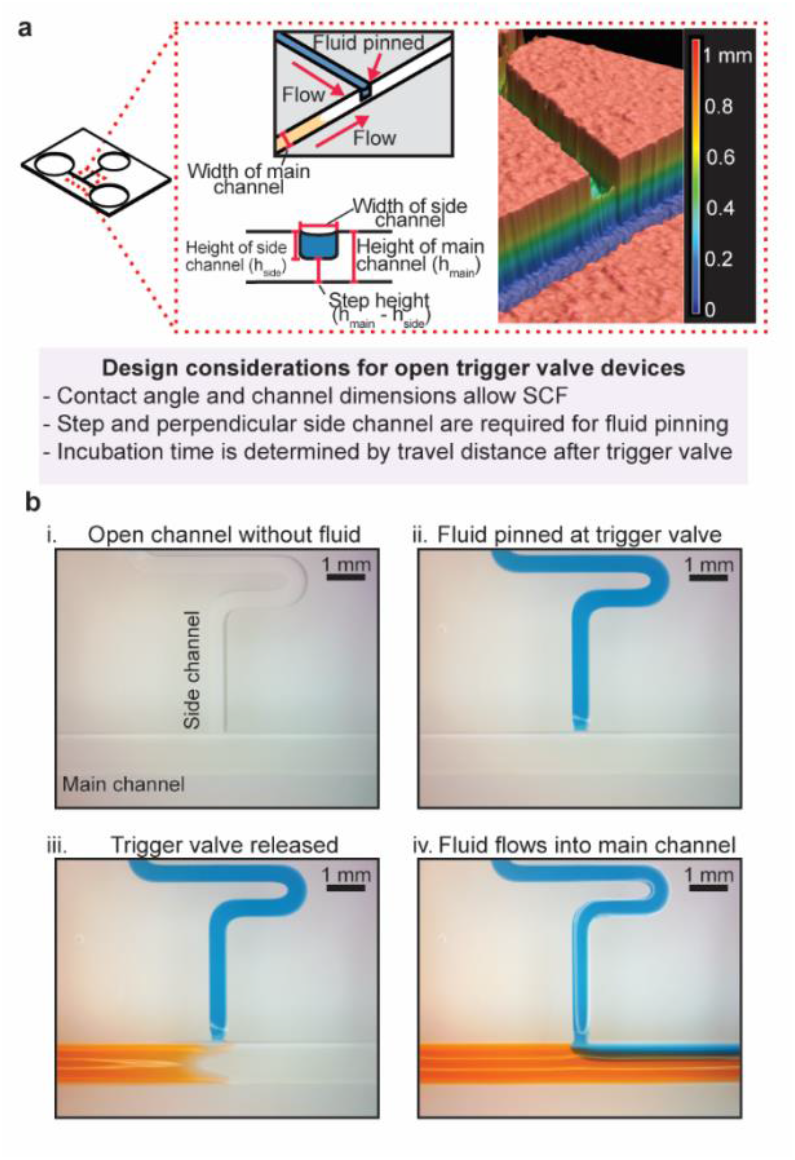
Considerations and operation of trigger valving in open channels. (a) Open-channel device with one TGV highlighting the intersection of the main channel and side channel and key aspects of TGV devices. (Inset) Drawings of the side channel opening indicating where pinning occurs (upper left panel) and a view of the trigger valve gate with the step from the bottom of the main channel to the bottom of the side channel (lower left panel)—a critical feature preventing flow into the empty main channel. Accompanying Keyence profilometer photo of an example open TGV (right panel). (b) A series of images illustrating TGV release in open channels starting with an empty main and side channel (bi) with the same channel dimensions as the sketch in i and ii. (bii) After liquid is added to the side channel, the fluid pins at the edge of the opening to the main channel where it remains stationary. (biii) When the main channel fluid makes contact with the pinned fluid, the side channel liquid is de-pinned. (biv.) Fluids from main channel and side channel flows then stabilize over time.

Open microfluidics has been used for many applications in organ-on-a-chip models, space science, neuronal networks, and cancer research.^27–30^ Open microfluidics leverages one or more air-liquid interfaces through the removal of one or more channel walls to enable improved accessibility of the channels. The popularity of open microfluidics is linked to several advantages such as accessibility, low cost, easy fabrication, and easy surface treatment.^1,4–7^ Through the removal of a channel wall, open channels afford the ability to add fluids at any point in a channel. Adding TGVs to open channels now add a layer of flexibility to autonomously introduce another flow of fluids to a channel – which opens the possibilities for hybrid open/closed devices. Many channel geometries, architectures, and valves reported in closed microfluidics may be applied to open microfluidic configurations to increase their capabilities and range of applications.

Drawing on prior work in closed-channel systems, we consider here the so-called two-layer or “stair-step” TGVs that have a higher stability than one-level valves due to the two-dimensional geometric expansion conferred by the separation of the main and side channels, which prevents leakage of pinned fluid (Figure 1a).^9,19,24^ In capillary-driven devices, the flow is regulated by surface energies and geometric features. Passive valves, which include TGVs, based on geometrical pinning are a common method for fine control of flow and on-chip programming. Geometrically, pinning occurs when capillary pressure is lost due to a sudden change in the channel architecture (such as an enlargement). Pinning of the fluid at the TGV “gate” is important to ensure that the fluid stops flowing until the depinning by a perpendicular flow by a miscible fluid. Retention of the liquid in the side channel until the main flow reaches the TGV relies on the liquid being pinned at the gate (or aperture), which refers to the edges of the side channel that intersects with the main channel (Figure 1b). Pinning of a liquid on edges depends on the edge angle and on the liquid-solid contact angle.^31–34^ If the constraints on the liquid (e.g., pressure) exceed the pinning angle threshold, pinning does not occur. Otherwise, pinning is stable, allowing us to leverage this phenomenon to create TGVs in open configurations.

In this work, we show that TGVs can be positioned in series in open-channel geometries through the de-pinning of multiple channels with immobilized fluids and that we can leverage characteristic features to design an open-TGV device. We also investigate the dynamics of flow and present an analytical model describing the travel distances and flow velocities based on the concept of friction length^3^ with a comparison to experimental results. Sequential de-pinning of trigger valves results in layered co-flows when the different liquids are miscible or have a very small liquid-liquid surface tension, enabling autonomous controlled fluid addition. The use of trigger valves was also used to demonstrate applicability through the detection of nitrites in water and meat samples.

## Results and Discussion

We present an analytical model of TGVs in open capillary channel configurations – a new tool for the open microfluidics tool kit. While the concept of open TGVs have been introduced in brief^26^, to our knowledge, this is the first in-depth development of theory complemented with a comparison to in-lab experiments. Throughout our study, three important characteristic aspects of the open TGVs were investigated: programmed TGV release, fluid velocity, and fluid layering.

### Part 1: Model

A sketch of an open capillary device with TGVs on the same side or opposite side of the main channel is shown in Figure 2a and b, respectively. Side channels are filled with immobile liquids (blue, yellow and purple). The “main” or “primary” fluid flows due to capillarity and sequentially opens the different gates, i.e., de-pins the immobilized side liquids which start co-flowing in the main channel. The side channels can be placed on the same side of the main channel (Figure 2a) or in a staggered arrangement (Figure 2b).

**Figure 2.**
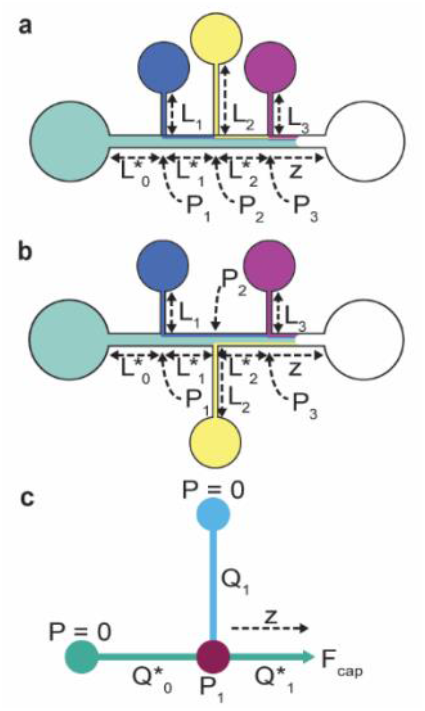
Sketch of the device with the three TGVs, placed on the same side (a) or on opposite sides (b), in the notations, a star denotes the main channel. (c): schematic of the flow at a node (intersection of the TGV and main channel) used for the calculation. The letter *Q* corresponds to a volumetric flow rate, and *P* to a pressure.

Figure 2c shows the principle of the calculation of the travel distance of the front meniscus (z) as a function of time after the front meniscus has passed a TGV. The flow is actuated by the capillary pressure at the front meniscus and balanced by the friction on the different wetted walls. In the inlet channel (before the first TGV),—and referenced by the index 0 in the calculation—the travel distance is determined by the generalized LWR (Lucas-Washburn-Rideal) law.^35^ Table 1 details symbols with their corresponding units.

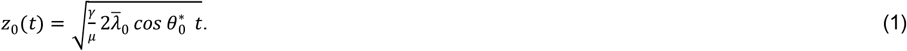

**Table 1:** Notations.

| Name | notation | unit | Remarks/References |
| --- | --- | --- | --- |
| Length of channel | $L_i$ | mm | Lengths of side channels |
| Length of main channel sections | $L_i^*$ | mm | Lengths between two nodes (TGVs) |
| Cross-sectional area | $S$ | mm<sup>2</sup> | Different for main and side channels |
| Average friction length | $\bar{\lambda}_i$ | mm | Characteristic of friction<sup>35</sup> |
| Perimeter (total) | $p_i$ | mm | Includes free perimeter |
| Perimeter (wetted) | $p_{w,i}$ | mm | Only wetted part |
| Perimeter (free) | $p_{f,i}$ | mm | Perimeter of free surface (in a cross-section) |
| Viscosity (dynamic) | $\mu$ | mPa.s | Nonanol |
| Surface tension | $\gamma$ | mN/m | Nonanol |
| Contact angle | $\theta$ | degrees | Static contact angle |
| Generalized Cassie angle | $\theta_{*i}$ | degrees | Generalized Cassie angle for open channels |
| Contact angle | $\theta_d$ | degrees | Dynamic contact angle |
| Pressure | $P$ | mPa | Pressure at the nodes |
| Capillary pressure | $P_{cap}$ | mPa | Laplace pressure of the meniscus |
| Flow rate | $Q$ | mm<sup>3</sup>/s | volumetric flow rate |
| Velocity | $V$ | mm/s | Averaged in a cross section (side channels) |
| Velocity | $V^*$ | mm/s | Averaged in a cross section (main channel) |
| Travel distance | $z$ | mm | Middle of advancing meniscus |
| Geometrical coefficient | $A_i$ | mm<sup>3</sup> | Side channel |
| Geometrical coefficient | $A_i^*$ | mm<sup>3</sup> | Main channel |
| Auxiliary geometrical coefficient | $B_i$ | mm<sup>3</sup> | Node coefficients |
| Time | $t$ | s | |

If necessary, relation (1) can be modified by a dynamic contact angle.^35,36^ The algebraic developments for the determination of flow past the TGVs are lengthy and are detailed in SI.1. We only present here the principle behind the model (Figure 2c). Let us introduce the geometric coefficients 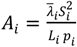 and 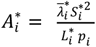 where the star (*) denotes the main channel and *i* is the channel index (Figure 2a). These coefficients have the dimension of volume (mm^3^) and solely depend on the width, depth, and length of each channel. The principle of the calculation can be easily shown considering the first TGV. The pressure at node 1, *P*_*1*_, is expressed by the two relations

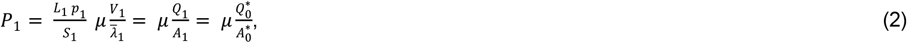

this can be combined with the mass conservation relation 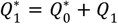 to produce the relation between the pressure at the node and the meniscus velocity 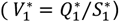:

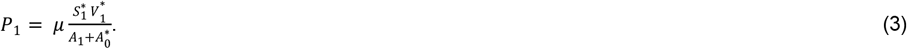

In the viscous regime, the capillary force balances the wall friction which, in terms of pressure, yields the relation

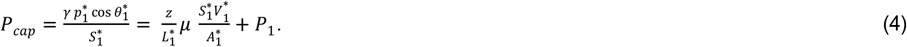

Using 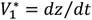 and combining (3) and (4) yields the solution,

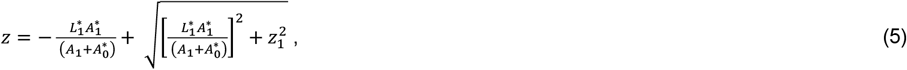

where *z*_*1*_ is the travel distance corresponding to the LWR law in the main channel, and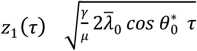, where *z*_*1*_ and *τ* are the distance and time starting from node 1. The principle can be extended by recurrence to an arbitrary number of TGVs as the side liquids are identical with the only difference being the slight changes from added coloring—and co-flow as a unique liquid downstream each trigger valve (see SI.1). We then obtain

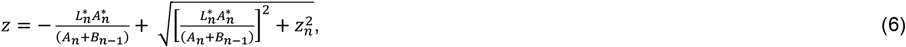

where the *B*_*i*_ are a mathematical sequence including all preceding 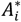 and *A*_*i*_.

### Part 2. Open flow experiments

#### 1. Programmed trigger valve release allows for tunable timing between fluidic events

We investigated how to achieve tunable timing between fluid events by using defined distances between TGVs. Towards this goal, we used the analytical model from Part 1 and validated the results through in-lab experiments (Figure 3). We designed two channels fabricated in poly(methyl)methacrylate (PMMA) with three TGVs (cross-section dimensions of w = 400 μm and h = 600 μm) oriented along the main channel (cross-section dimensions of w = 1 mm and h = 1 mm). The TGVs in Devices 1 and 2 were separated by travel distances of 15 and 30 mm, respectively. Nonanol was chosen as the main and side channel fluid with slight differences in coloring to visualize the flow, but constant viscosity. To ensure sustained fluid pumping, a paper pad was added to the end of the main channel. For Device 1, the time between the release of TGV one and two was 3.70 (±0.05) seconds and the time between TGV release of two and three was 3.93 (±0.06) seconds. For Device 2, the time between TGV release increased with the longer channel length to 7.67 (± 0.10) and 9.83 (±0.20), respectively. (Figure 3a and b, designs in Figure SI.4.1). Data reported are the average of three trials with the standard deviation. Data for all trials can be found in Figure SI.5.1. Comparison of the experimental data to the theoretical model for fluid front travel distance and velocity, correspond closely to each other, validating that our model predicts fluid flow in open channels with integrated TGVs.

**Figure 3.**
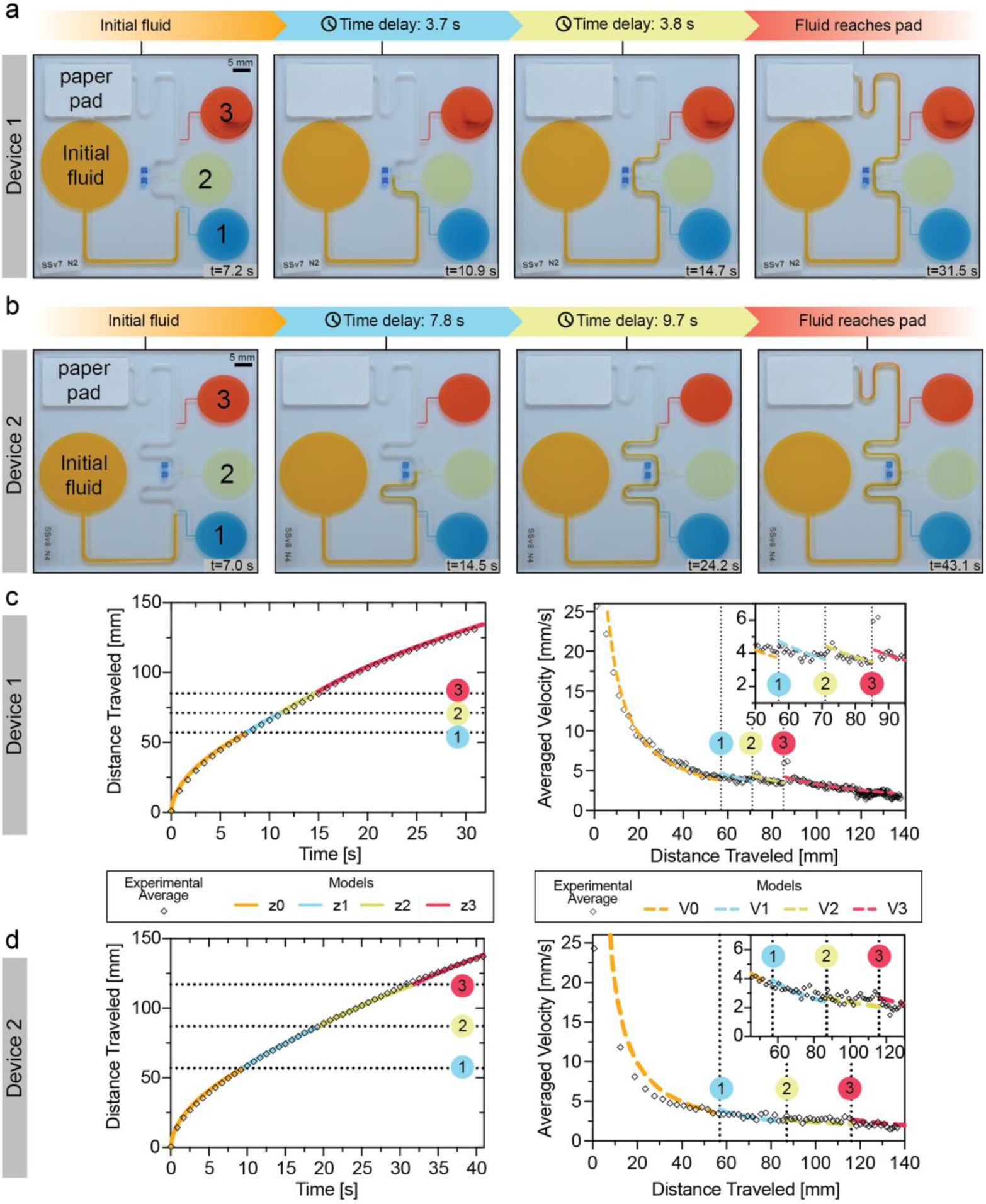
Distance between TGVs provides timed control of fluid release. Progression of flow in devices with parallel TGVs spaced approximately 15 mm (a, device 1) and 30 mm (b, device 2) apart. Comparison of model (solid line) with experimental fluid front travel distance (diamonds) at the meniscus over time for device 1 (c, left) and 2 (d, left). Experimental data was averaged for three trials (n=3). Model is presented in segments corresponding to the calculated travel distances prior to the TGV release (orange, z0), between the first and second TGV (blue, z1), between the second the third valves (yellow, z2) and between the third valve and the paper pad (red, z3). Experimental velocities (diamonds) were calculated from the travel distance and fluid velocities of each channel were calculated from the travel distance progression and are compared against the calculated model velocities (dashed line) using a dynamic contact angle model (V0, orange) for the inlet. Model is shown at the first (blue, V1), second (yellow, V2), and third (red, V3) valves due to the increase in velocity upon TGV release. Data is averaged from three (n=3) trials. Raw data can be found in Figure SI.5.1.

Like closed channels, implementing TGVs in open-channels is important for on-chip control of fluid addition and programming of timed incubations, waiting periods, or wash steps for the application of this tool in assays, workflows, and reactions such as antibody patterning, polymerase chain reactions (PCR) or enzyme-linked immunosorbent assays (ELISA). Channel geometries and side channel flow rates can be designed to control how much fluid is added to the channel to achieve a certain final concentration or add a set amount of fluid into the flow. Moreover, the ability to implement this method of fluid addition in open channels could enable the automation of fluid addition by controlling when a fluid or reagent is added and the speed/flow rate at which the fluid is introduced, thus reducing the need for additional manual pipetting.

#### 2. Investigation of dynamics in open trigger valves

We further investigated the effect of TGV release on the main channel fluid dynamics. For this study, we used parallel TGVs spaced 3.70 mm apart (Images of flow progression can be found in Figure 4a). Using nonanol, we measured the travel distance over time (Figure 4e and f) and noted the spatial locations of the TGVs (black dotted lines). Upon closer inspection of the TGV region of the graphs (Figure 4c, inset), the model predicts a stair-like trend where the TGVs cause an increase in fluid velocity. Shortening the distances between the valves makes this trend more evident (Figure 4 c and d). The release of a TGV increases the flow velocity where the side channel acts as an additional reservoir closer to the front meniscus than the inlet reservoir on the main channel. Therefore, the velocity increases in the main channel at the opening of the TGV. Notably, the experimental data shows correlation with these predicted jumps. A comparison of experimental results to the theory for TGVs on opposing sides can be found in Figure SI.5.2 and raw data for valves on the same side and on opposite sides can be found in Figure SI.5.3. Further, an investigation of an increased travel distance between the last trigger valve and the paper pad showed the model is valid for even longer channel lengths (Figure SI.5.4)

**Figure 4.**
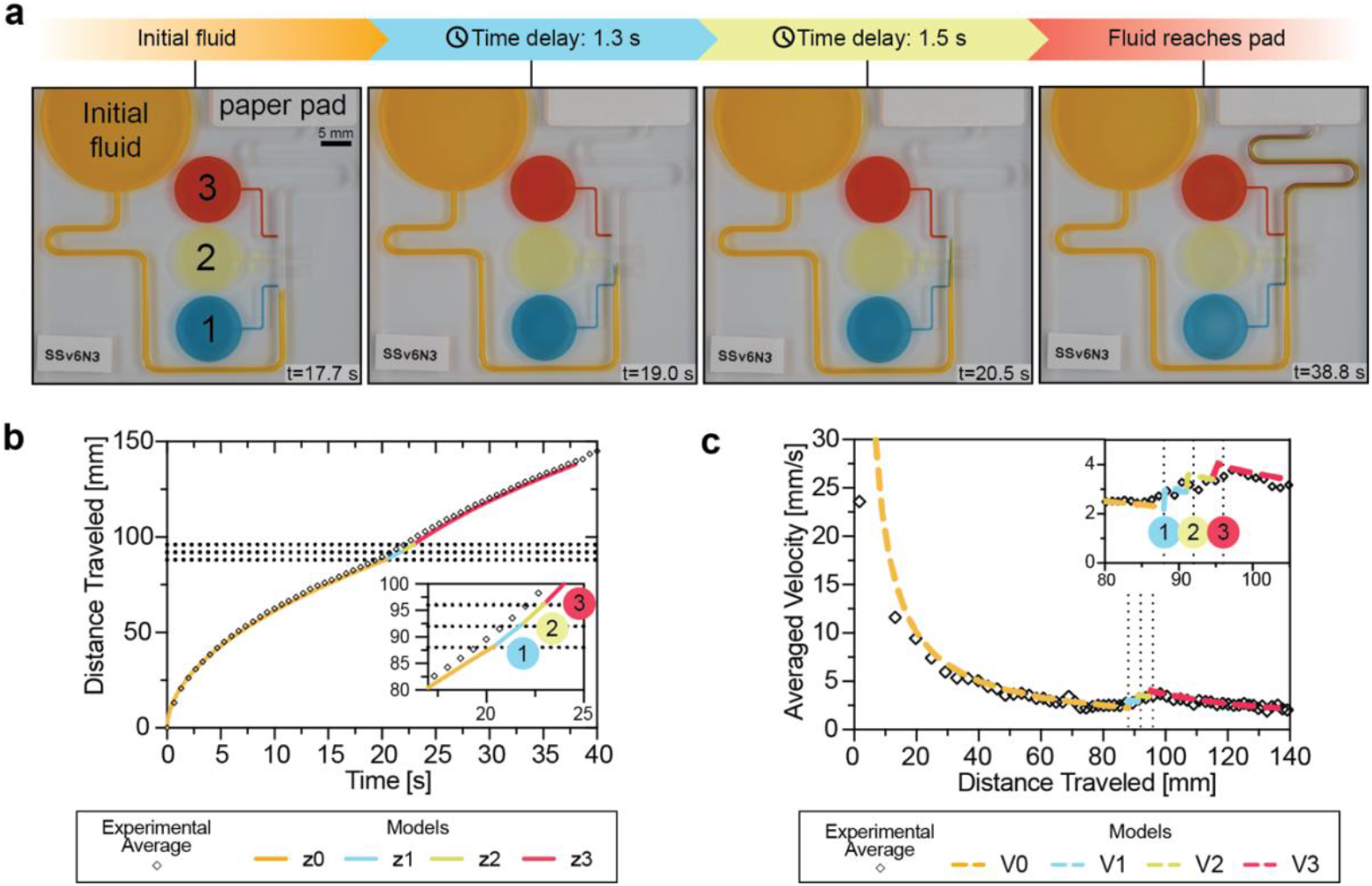
Open channel flow experiments show an increase in velocity after each valve opening. Progression of fluid flow through device with shorter distances between TGVs (a). Comparison of the theoretical model (solid line) with experimental fluid front travel distance at the meniscus (diamonds) for devices with three side channels in parallel. Experimental data were averaged for four trials (n=4). Model is presented in segments corresponding to the calculated travel distances prior to the TGV release (orange, z0), between the first and second TGV (blue, z1), between the second the third valves (yellow, z2) and between the third valve and the paper pad (red, z3). Fluid velocities were calculated from the travel distance and the experimental data (diamonds) compared against the calculated model (dashed line) velocities using a dynamic contact angle model (orange, V0) for the inlet. Model is shown at the first (blue, V1), second (yellow, V2), and third (red, V3) valves due to the increase in velocity upon TGV release. Raw data can be found in Figure SI.5.3.

In a comprehensive view, the release of the fluid at a TGV increases the velocity of the main channel in the geometrical configuration of the present device; however, in reality, this effect is more complex. The lengths and widths of the side channels as well as the distances between the valves are determining parameters for the velocity jump in the main channel at a TGV (these dimensions are included in the constants *B*_*i*_ and *A*_*i*_ of the model). More specifically, the wall friction in the side channels conditions the velocity jump. A detailed analysis is done in SI.2. Briefly, there is a positive velocity jump at a TGV, and this jump is higher when the friction in the side channel is less. Thus, when we consider this in an applications view, the lengths of the side channels, distances between the side channels, and the cross-section dimensions are critical aspects to take into consideration. In cases where specific conditions need to be met, the presented model allows users to predict the fluid velocities and flow rates of the side channel pumping which can inform users in determining optimal dimensions.

#### 3. Trigger valves enable layering in open systems

We investigated the co-flowing of fluids after the release of each TGV. With the pinning of each fluid at the gate and the subsequent release by the main channel fluid, each valve demonstrates a “burst” phase into the main channel where the fluid starts flowing into the channel. The thickness of the layer is initially large, but stabilizes over time. Using dyed nonanol and devices with parallel TGVs (Figure 4), layer width measurements were taken at 0.5 mm after the valve in the main fluid channel for each of the valves, monitoring only the topmost fluid layer. The measurements were taken until the fluid exited the frame. Figure 5 shows the layering and a comparison of the layer thicknesses over time and a side view image of the fluid layers for trigger valves on the same side of the main channel. Layer thickness comparison for valves on opposing sides of the main channel can be found in Figure SI.5.6.

**Figure 5.**
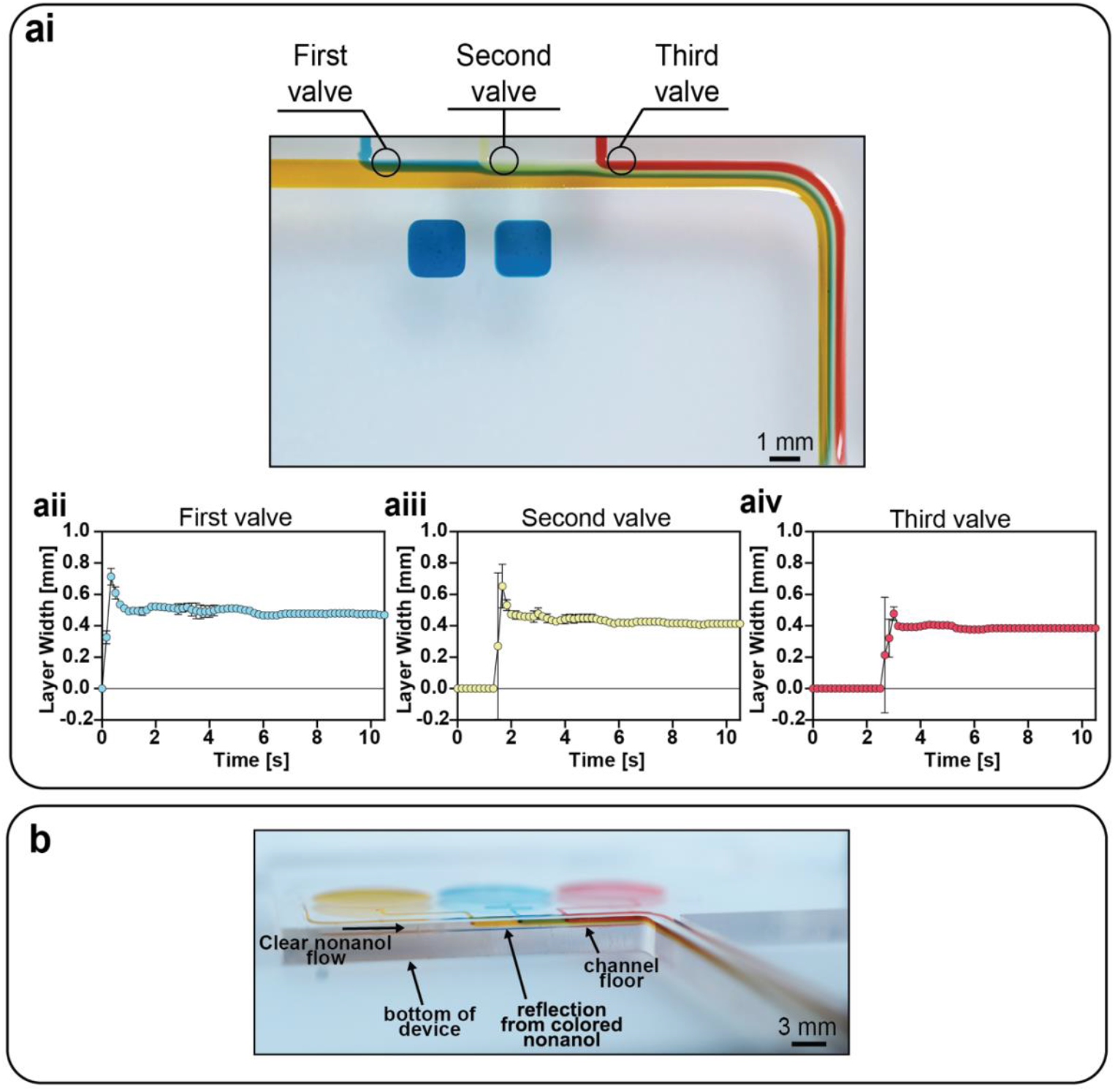
Fluid layers released by the TGVs stabilize in width over time. (ai) Image of the fluid layers in a parallel configuration. Circles indicate where layer thickness measurements were taken for each layer (0.5 mm after each valve). Plots of the measured layer widths over time for the parallel channel configuration after the first (aii), second (aiii), and third (aiv) valves. Data are shown as an average of three replicates (n = 3) with error bars indicating standard deviation. Layer widths are reported at a defined point in the channel (see diagram in SI.5.7). A side view image of the fluid layering after all three valves were released by undyed (clear) nonanol. Order of colored nonanol used in the side channels for part b were adjusted for visualization of the layers and are not the same as in part a.

**Figure 6.**
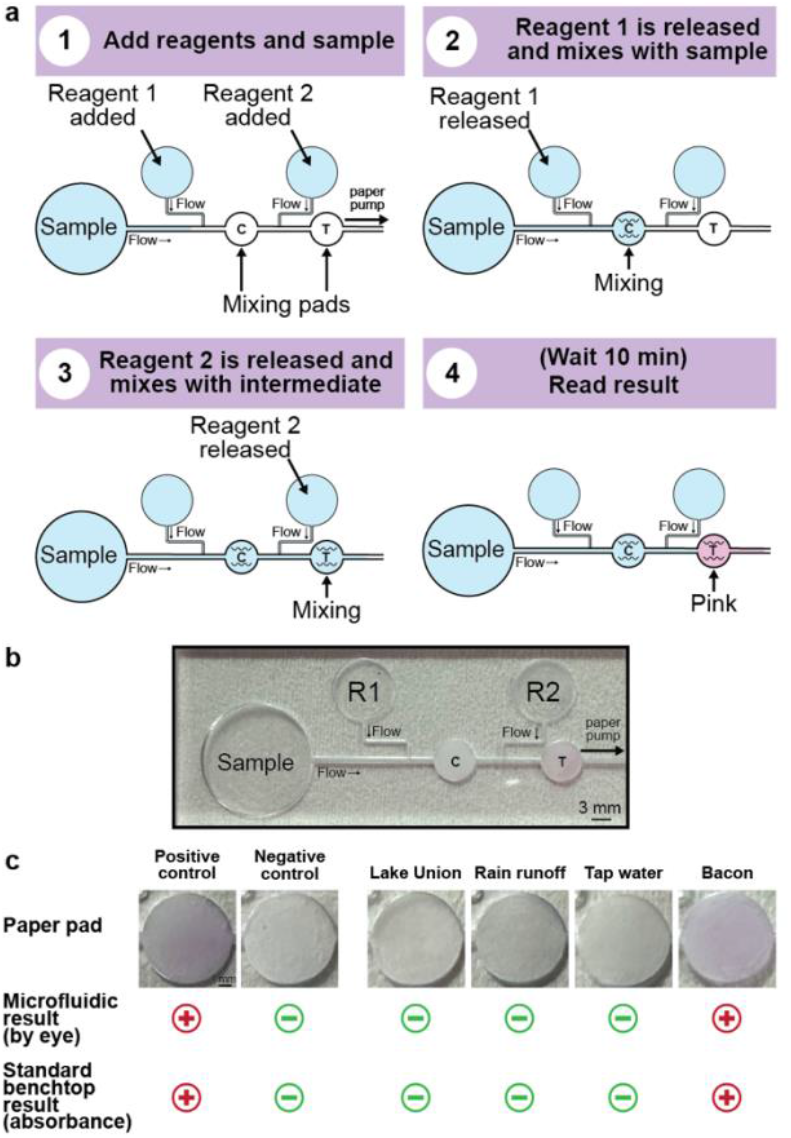
Trigger valves enable on-chip nitrite detection via the Griess reaction. (a) Schematic of device operation using the Griess reaction. Reagents 1 and 2 (R1 and R2) are added to the side reservoirs and are subsequently released by the flow of the sample with interspersed mixing steps via circular paper pads. Pads function as mixers as well as control (C) and test (T) zones. (b) Representative photograph of a device to detect nitrites in bacon samples. (c) Representative images of paper pads from Griess reaction with a 5 mg/L nitrite ion standard solution (positive control), deionized and distilled water (negative control), Lake Union (Seattle, Washington) water sample, rain runoff water sample, Seattle tap water sample, and a bacon sample. A red plus sign denotes a positive test result; A green minus sign denotes a negative test result (pink not visible by eye or spectrophotometer readout yielding a concentration below 1 mg/L). Experiments were done in triplicate (n=3) with different devices (Figure SI.5.9). Concentration data for the environmental and meat samples measured using the standard benchtop method can be found in Table SI.5.1.

After each TGV release, we observed an increase in layer width at the initial opening and then a stabilization of the fluid layers over time with uniform layering of the topmost layers. Moreover, the fluid layers become a cascade of layer thickness with the succession of TGV opening in series where the topmost layer (layer closest to the channel wall) is wide and previously released layers become thinner (Figure 5aii-aiv). The decrease in width is observed to be due to the previous layers shifting downwards into the main channel and the most recently released layer on top demonstrating an observed 3D effect– a complex phenomenon for the work of a future study. Despite this effect and the initial burst at the gate, layering is uniform between each valve as seen in the Figure 5ai to aiv and we have maintained separation of the layers through 30 mm after all the TGVs are released with the continued capillary pumping by the paper pad. Additional layering data for opposite-facing trigger valves can be found in Figure SI.5.6. A diagram of the location of measurement with respect to each valve can be found in Figure SI.5.7. True layering measurements in 2D and 3D and a detailed study of the underlying fundamental fluid dynamics are focal points for future work.

Understanding the effect of fluid layering is an interesting aspect of fundamental open microfluidics. Layering or laminar co-flows are important in the applications of TGVs to create functional devices as this can be used to create a liquid-liquid interface for chemical reactions or on-chip patterning of hydrogels and antibodies. Comparatively, in closed channels, similar flows have been achieved; however, our system enables access to the channel contents for manipulation or interaction with the flows if needed for a specific application. Open-channel devices are also well suited for studies of the mixing of the channel contents, although fluid mixing is not demonstrated here as shown in Figure 5a.

The reason for this absence of mixing stems from the smallness of the molecular diffusion compared to the axial convection. Symbols with the corresponding definition and units are presented in Table 2. The Taylor-Aris dispersion theory compares the axial convection time scale *t*_*conv*_∼ L/*V* to the radial diffusion time *t*_*diff*_∼ *δ*^2^/*D*. Because the diffusion coefficient *D* in liquids is small, *t*_*diff*_ is large and the length where the two characteristic times balances L is given by^37,38^

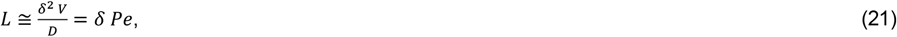

where *δ* is a typical dimension of the fluid layer (the width in our case), *V* is the liquid velocity, *D* is the molecular diffusion coefficient and *Pe* is the Peclet number (*Pe* =*δV*/*D*). The diffusion coefficient of nonanol can be approximated by the Stokes–Einstein–Sutherland equation, for diffusion of spherical particles through a liquid with low Reynolds number

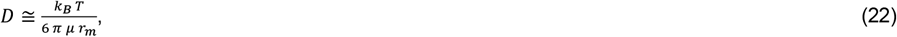

where *r*_*m*_ is the molecular radius, *T* is the temperature and *K*_*B*_ is the Boltzmann constant. Per literature^36^, the nonanol radius was found to be of the order of 0.4 nm and the constant *k*_*B*_ *T* is approximately 4.11 pN.nm at room temperature. Using the value of 0.011 Pa.s for the viscosity of nonanol, we obtain *D*∼ 4 10^-11^ m^2^/s. If we consider a liquid “filament” or “layer” coflowing with the carrier flow, with a typical width of 200 µm, the Taylor-Aris diffusion length is of the order of *L* ∼ 50 –100 cm for a velocity on the order of 1 – 2mm/s. In this case, the molecular diffusion does not play an important role in the flow downstream of the TGVs along a length of 20 – 50 cm. The convective velocity *V* of 1 mm/s is much larger than the radial propagation *D*/*δ* of 0.25 µm/s: considering a duration of 50 seconds, molecules in the layer have moved by 50 mm in the axial direction while the radial diffusion is only 12.5 µm.

**Table 2:** Notations.

| Name | Notation | Unit | Remarks/References |
| --- | --- | --- | --- |
| Time | $t$ | s | |
| Mixing Length | $L$ | mm | Length where noticeable mixing occurs |
| Hydraulic radius | $R_H$ | mm | |
| Peclet number | $Pe$ | Non-dimensional | |
| Viscosity | $V$ | Pa.s | Fluid's resistance to flow |
| Molecular diffusion coefficient | $D$ | m<sup>2</sup>/s | |
| Boltzmann constant | $k_B$ | J/K | |
| Temperature | $T$ | K | Kelvin |
| Molecular radius | $R$ | pm | |
| Reynold's number | $Re$ | Na | |
| Dean number | $De$ | Na | |

However, the situation would be different when using different liquids and/or narrower channels. For example, the coefficient of diffusion of water is fifteen times higher than nonanol (*D*∼10^-9^ m^2^/s) and even if the velocity of the flow is twice that of nonanol, we find Taylor-Aris dispersion lengths of the order of 20 cm. The same reasoning suggests that there is probably also a lower limit to the size of the layers: in the case of nonanol, the Taylor-Aris dispersion length decreases to 25 cm for a layer width of 100 µm. Hence, a layering process is not automatically obtained.

On the other hand, there are cases where mixing is desirable, such as fertility applications where one fluid (purified sperm preparation) needs to be mixed with another (sperm cryoprotectant) at a planned rate. In this case, mixing features—inspired by the many devices developed for forced flows— can be added in the main channel after the last TGV. For example, turns might promote mixing. This effect— called Dean effect—is due to the formation of a vortex in the turn. The formation of a vortex in a turn is characterized by the Dean number^39,40^

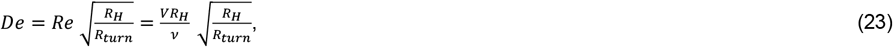

where *De* is the Dean number and *Re* the Reynolds number. The Dean number must be larger than 1 to observe a vortex in the turn. In our case, *V* ∼ 3 mm/s, *R*_*H*_ ∼1 mm, *v*∼1.3 10^-5^ m^2^/s, so that *Re*∼0.2 and *De*∼0.1 which explains the absence of mixing in the turns.^39,40^ Note that this low Dean number is due to the high value of the viscosity and relatively low surface tension of nonanol that condition the velocity of the open capillary flow. Complete mixing is also not seen in the paper pads as shown by the flow profile of the paper pads (Figure SI.5.8), but a deeper study in the mixing within the paper pads is a focus for future work. However, this might not be the case for “faster” liquids, such as water, isopropyl alcohol (IPA) solutions, etc. A definition of “faster” liquids is given in reference^36^.

Lastly, coloring the different layers of the flowing liquids enables an accurate observation of the open capillary flow in a turn. It is observed that the innermost part of the fluid flows first in a turn, then the outer fluid catches up. This observation has already been reported in the literature^41^ which is due to the travel length of each layer remaining the same.

#### 4. Trigger valves enable nitrite detection in open channels

Lastly, as a proof-of-concept, we applied the open channel trigger valves to the detection of nitrites in water and meat samples. This approach takes advantage of the precise control of capillary flow dynamics to facilitate sequential reagent flow and mixing. For nitrite detection, the Griess reaction was exploited using sulfanilamide (Reagent 1) as aromatic amine and N-1-(naphthyl)ethylenediamine (Reagent 2) as a coupling agent from a commercially-available nitrite colorimetric test kit. The Griess reaction is an azo coupling mechanism, where an aromatic amine in acidic conditions reacts with nitrite ions to form a reactive diazonium salt, which can be combined with an electron-rich coupling agent to form a diazo dye characterized by absorption of visible light at 540 nm. This reaction enables a colorimetric response visible by the naked eye with a pink color, the intensity of which correlates with nitrite concentration. By utilizing open channel TGVs, the system ensures controlled successive interactions between reagents and the analyte; it is important that the sample is first mixed with the Reagent I before mixing with Reagent 2. We designed a device (engineering drawing found in Figure SI.4.2) with two TGVs and two circular mixing paper pads (one after each TGV) to facilitate reagent mixing. Reagent 1 is released by the first valve and flows into the main channel to reach the first mixing paper pad together with the sample. The presence of a paper pad contributes to mixing the reagents and forming the diazonium salt intermediate before reaching the second valve. When the mixing flow reaches the last valve, reagent 2 is released and mixed in a second circular paper pad, enabling the final stage of mixing. This second pad also functions as the test pad for result evaluation by the naked eye through a change in color from white to pink if nitrite ions are present.

Using our device, we tested deionized and distilled water and a 5 mg/L nitrite ion standard solution as negative and positive controls, respectively. We expected no color change in the deionized and distilled water and a pink color change in the nitrite standard solutions, which were observed in our experiments. We then applied this work to environmental samples of water collected from Lake Union (Seattle, Washington), rain runoff from a rose garden, and a Seattle water tap. As nitrite can be found in fertilizers and agricultural lands, we chose various water sources to test for nitrite. In these tests, we did not detect a color change by eye. We then compared our results to the gold standard benchtop method through absorbance measurements; the absorbance values were below our set threshold of 1 mg/L (the United States Environmental Protection Agency (EPA) Maximum Contaminant Level (MCL) for drinking water). Lastly, we tested commercially-available bacon strips for the presence of nitrite, a commonly used food additive. We were able to detect visually the presence of nitrite via our microfluidic device and confirmed this result through the gold standard benchtop method. These systems are particularly suited for point-of-need applications in environmental monitoring, where rapid and reliable detection of nitrite and other analytes is critical. Taken together, we conclude that open trigger valves can be used to simplify workflows for multistep reactions.

## Conclusion

In this work, we have developed the concept of a TGV for capillary flows in open channels, building on prior work on valving in closed channels^8–10,12–24^. We demonstrate that the TGV principle is compatible with open capillary flows using the example of stair-step TGVs. This type of valve is highly effective because the side fluids are stably immobilized by the sharp solid edges on three sides of the aperture and the presence of air on the fourth, upper side. This work—while fundamental here—has the potential to be useful in a multitude of fields such as biomedical research, biosensors, and point-of-care diagnostics.

Using TGVs in series—on the same side or on opposite sides of the main channel—results in the multi-layering of fluid flows. In conventional microfluidics, co-flows have already been achieved due to the high degree of laminarity. Here, we show that layered co-flows can also be achieved for capillary flows in open channels. This principle is also valid for closed-channel capillary flows. The layering continues along the entire channel due to poor molecular mixing caused by low Reynolds numbers. This occurs when the channel length is much shorter than the threshold length predicted by the Taylor-Aris dispersion theory. At low velocities (approximately <5 mm/s) and intermediate to high viscosities (>4 mPa.s; e.g., nonanol capillary flow)^36^, no vortex forms in the channel turns—in line with Dean’s vortex theory—and no liquid mixing occurs. However, this applies to very viscous liquids. Analyzing Dean’s flow in turning open channels is a logical extension of this work. In addition, we observed a 3D effect in the co-flows where the fluid layers overlap along the flow, which will be a focus of future work.

The flow dynamics can be predicted by an analytical model based on the fundamental concept of average friction length, which describes wall friction. In the viscous mode—quickly established in the channels—the model balances the constant capillary force (as long as the main channel maintains a constant cross-section) and the pressure drop due to friction in the main and various side channels. A recurrence relation is derived to calculate the flow dynamics for an arbitrary number of TGVs.

Very narrow co-flows are of great interest for applications in biology and chemistry. This work shows that very narrow liquid layers can co-flow in an open channel depending on the geometric configuration of the channels. For example, with three TGVs in series, the width of the fluid layers is less than one-fourth of the main channel width. Achieving smaller scale widths is important for open microfluidic applications and opens a new avenue for small scale patterning of fluids (e.g., hydrogels^42^), chemical micro-reactions, or could be a precursor to mixing.

## Methods

### Device design and fabrication

Two general device configurations were designed for this study: three TGVs oriented along the same side and the opposite side of the main channel. All devices had a main channel with cross-section dimensions of width and height equal to 1 mm. The side channels had cross-section dimensions of width = 400 µm, height = 600 µm with a TGV gate at the intersection of the end of the side channel and the main channel. Each device differed by the total length of the main and side channels, distance between TGV gates, and the configuration of the side channels along the main channel. For the Griess reaction device, two trigger valves were used and a circular well (7.1 mm diameter) were added after each trigger valve for reagent mixing. Images and schematics of two representative designs can be found in Figure 3. Engineering drawings of all the devices used in this study can be found in Figure SI.4.1 and SI.4.2.

Computer-aided designs (CAD) and computer-aided machining (CAM) G-code (.simpl) files for the devices were created in Autodesk Fusion (Autodesk, San Francisco, California). Devices were milled using a Datron Neo computer numerical control (CNC) mill (Datron Dynamics, Milford, New Hampshire) in 3.175 mm poly(methyl)methacrylate plates (#8560K239; McMaster-Carr, Sante Fe Springs, California) for the open-channel flow dynamics experiments or 4.00 mm polystyrene plates (#ST31-SH-000200; Goodfellow Corporation, Pittsburgh, Pennsylvania) for the nitrite detection via the Griess reaction. The channels were milled to have rounded inner corners to prevent the formation of capillary filaments using endmills with a cutter diameter of 1/32” (TR-2-0312-BN) or 1/64” (TR-2-0150-BN) from Performance Micro Tool (Janesville, Wisconsin). Channel dimensions and the quality of the milled cuts were verified using a Keyence wide area 3D measurement VR-5000 profilometer (Keyence Corporation of America, Itasca, Illinois). The channel bottom is estimated to have a few microns of roughness due to the Datron milling process, which is one magnitude below the roughness values observed by Lade *et al*. to produce substantial fluctuations in velocities in the capillary flow.^43^

After fabrication, the devices were ultra-sonicated in 70% (v/v) aqueous ethanol for 30 minutes to remove any residues and debris using the Branson M2800H. The cleaning solvent was reused no more than five times. After sonication, the devices were rinsed in fresh 70% (v/v) aqueous ethanol and subsequently in deionized (DI) water. Devices were then added to a bioassay dish and partially covered with the lid to dry in the hood overnight. For the devices milled in polystyrene, devices were oxygen-plasma treated with a Diener Zepto PC EX Type PB plasma treater.

Paper pads for the devices were designed in Adobe Illustrator 2023 (Adobe, San Jose, California) and cut into 15.2 mm wide and 25 mm long rectangular pieces or into a circular shape with a 7.00 mm diameter. The paper pads were Cytiva Whatman #1 filter papers (#1001-185) and cut out using the Graphtec CE-7000 plotter cutter and the Cutting Master 5 program (Graphtec America, Irvine, California). Paper pads were stored in a bioassay dish prior to use.

### Solvent preparation and physical properties

For trials conducted with channels on the same side in our flow velocity experiments, nonanol (Sigma-Aldrich, # 131210) served as the flowing liquid. Dyed nonanol solutions were made using Sudan I (Sigma-Aldrich, # 103624), Sudan III (Sigma-Aldrich, # S4131), Solvent Green 3 (Sigma-Aldrich, #211982), and Solvent Yellow 7 (#S4016), each at concentrations of 0.5 mg/ml or 1.43 mg/ml. For all other devices, stock solutions of dyed nonanol were prepared using Sudan I, Sudan III, Solvent Green 3, and Solvent Yellow 7 in 10 ml volumetric flasks at concentrations of 1 mg/ml. These solutions were subsequently diluted with nonanol to concentrations of 0.5 mg/ml or 1.43 mg/ml for all trials.

For the Griess reaction, sodium nitrite (#S2252, Sigma-Aldrich, St. Louis, Missouri) in H_2_O was used as flowing liquid. A stock solution of 750 mg/L of sodium nitrite in ASTM H_2_O (#6442-85, Harleco, was diluted up to reach a concentration of nitrite ions of 5.0 mg/L. Griess reactants, sulfanilamide (Reagent I) and N-(1-Naphthyl)-Ethylenediamine in hydrochloric acid (Reagent II) from the “Nitrite/Nitrate Colorimetric Test” (Roche) kit (#11746081001, Millipore Sigma-Aldrich, St. Louis, Missouri). Reagents I and II were used as provided and not diluted. Environmental water samples were collected and stored at 4°C prior to sample preparation as followed: a 150 mL water sample was collected from Lake Union, Seattle, Washington with a Corning polystyrene bottle, a 15 mL sample of rain runoff from a rose garden was collected with an eyedropper into a 15 mL Falcon conical tube, and commercially available bacon (Private Collection center-cut double smoked bacon) was procured at a local grocery store. Tap water was collected in a 15 mL Falcon conical tube prior to experimentation. The water sample collected from Lake Union and the rain run-off were filtered twice using a 0.45 µm nylon (Fisherbrand Basix, Thermo Fisher Scientific, Waltham, Massachusetts) and a 0.2 µm Whatman cellulose acetate (CA) syringe filter (#6901-2502, Cytiva, Danaher Corporation, Washington, D.C.) to filter out any sediment and debris. The bacon was processed as described by Crowe *et al*.^44^ The bacon sample was then filtered three times using a 190 µm fine mesh paint strainer (Magca LTD), 0.22 µm CA vacuum filter (Corning Incorporated, Corning, New York), and lastly, a 0.2 µm CA syringe filter.

### Open-channel flow experiments

The devices were positions on a white background atop an adjustable lab jack. Videos capturing the progression of solvent flow in the devices were recorded using a Nikon D5300 ultra-high resolution single lens reflective (SLR) camera at 60 fps. Tabletop photography lights were adjusted to reduce shadows in the video. To obtain the distance that the fluid in the main channel traveled over time and/or to demonstrate fluid layering, the dyed nonanol was used and a paper pad was inserted into the rectangular outlet reservoir.

For the devices with extended lengths (Figure 3) between the TGVs, 106.02 µL of blue, yellow, and red dyed nonanol were added to the first, second, and third side channels, respectively. For the devices with 3.70 mm between the valves (Figure 4 and 5) on the same side and opposite sides of the channel, 57.02 µL were used instead. A visual check for depinning at the gate was done to ensure fluid flow is stopped. Afterwards, 490 µL of orange-dyed nonanol was added to the circular main reservoir. Devices were left to flow to release the TGVs and data collection stopped when the paper pad wicked the fluid. For the travel distance and fluid velocity experiments, data collection stopped when the paper pad wicked the fluid. For the layering experiments, the device with the valves on the same side followed the same order of colored nonanol in the side channels as the fluid velocity experiments. The device with TGVs on opposite sides of the main channel, blue, red, and yellow dyed nonanol were added first, second, and third TGVs, respectively, for enhanced contrast between the layers and better visualization of the fluid layering.

### Nitrite detection via the Griess reaction experiment

A calibration curve was generated using a 96-well plate and the nitrite detection kit with standards (in triplicate) of the following concentrations: 5.00, 3.75, 2.5, 1.00, 0.5, 0.25, 0.1, and 0.05 mg/L (0.05 mg/L sample as not included in the calibration curve due to a negative absorbance value). Water samples from Lake Union, rain run-off, faucet tap, and bacon were prepared in triplicate wells and prepared as described by the kit protocol. After 15 minutes, the absorbance of each well at 540 nm was measured using a Cytation 5 cell imaging multimode reader (Agilent Biotek, Santa Clara, California). The mean absorbance values were used for the calibration curve and the samples. Sample concentrations were interpolated using the equation of the standard curved and reported as a mean concentration (± standard deviation) (Table SI.5.1).

For the microfluidic experiment, two circular paper pads on top of each other were inserted into the circular wells and one rectangular paper pad was added to the end reservoir. 57.02 µL of Reagent I and Reagent II were added to the first and second side channels, respectively, into the device shown in Figure SI.5.9. 300 µL of nitrite sample were added to the main reservoir and devices were left to flow to release the TGVs and mix the reactants. Six refills of 57.2 µL were done in sequence to maintain Laplace pressure. In the future, the device geometry could be modified to eliminate the need for refills. Fluids were left to flow for 10 minutes after the release of the second valve. At the 10-minute mark, the device was flipped over onto a TechniCloth (#TX609, TexWipe, Kernersville, North Carolina) to stop the flow. On the backside, the second paper pad was visually evaluated for a pink color change by comparing the color with the color of the control paper pad (the first paper pad along the main channel). Images were taken with an iPhone 14 Pro smartphone camera.

### Image capture and data analysis

Images for the fluid dynamics were analyzed as described by Tokihiro *et al*.^36^ In brief, an image was captured every 10-30 frames using a custom Python program for the travel distance and velocity analysis. From the images, the fluid front was tracked using the segmented line and measure tools in ImageJ (National Institutes of Health). The resulting measurements were exported as a .csv file. For the layering images, every frame was extracted from the video using the custom python program for the layering measurements. The thickness of the topmost layers was measured after 0.5 mm from each valve until the fluid front reached the end of the frame. The measurements were exported as a .csv file. MATLAB codes were prepared for theory projects and are included in the supplementary material.

## Acknowledgements

Research reported in this publication was supported by the National Institutes of Health National Institute of General Medical Sciences grant R35GM128648 (A.B.T.), National Center For Advancing Translational Sciences grant TL1TR002318 (J.C.T.) and KL2TR002317 (T.M.N.), and the University of Washington. D.G. thanks Ph.D. programs on green topics (PON “Research and Innovation” 2014–2020) funded by FSE REACT-EU. We also acknowledge the M.J. Murdock Diagnostics Foundry for Translational Research. The content is solely the responsibility of the authors and does not necessarily represent the official views of the National Institutes of Health.

## Author contributions

J.C.T., A.B.T., J.B., E.B. designed the study. A.B.T., J.B., and E.B. are the principal investigators. J.C.T., I.H.R., D.G., and J.B. drafted the manuscript. J.C.T., I.H.R., D.G., and A.S. performed the in-lab experiments. J.C.T., I.J., D.G., and J.B. conducted the data analysis. J.B. developed the mathematical model and interpreted the data with J.C.T. The following provided high-level advising, conceptualization of the work, and manuscript revisions: E.M., T.M.N., A.O., A.B.T., J.B., and E.B. All authors read and approved the final version of the manuscript.

## Declaration of competing interests

A.B.T. reports filing multiple patents through the University of Washington and A.B.T. received a gift to support research outside the submitted work from Ionis Pharmaceuticals. E.B. is an inventor on multiple patents filed by Tasso, Inc., the University of Washington, and the University of Wisconsin-Madison. T.M.N. has ownership in Tasso, Inc.; E.B. has ownership in Tasso, Inc., Salus Discovery, LLC, and Seabright, LLC and is employed by Tasso, Inc.; and A.B.T. has ownership in Seabright, LLC; however, this research is not related to these companies. The terms of this arrangement have been reviewed and approved by the University of Washington in accordance with its policies governing outside work and financial conflicts of interest in research. The other authors declare that they have no known competing financial interests or personal relationships that could have appeared to influence the work reported in this paper.

## Availability of materials and data

All data generated or analysed during this study are included in this published article (and its Supplementary Information files).

## Notes

### Summary of Updates

Manuscript edited for minor edits to introduction and conclusion. Addition of Figure 6 and new seciton on nitrite detection in Results and Discussion. Author order updated. Supporting information updated.

