## Supporting Information for "The dynamics of capillary flow in an open-channel system featuring trigger valves"

#### Table of Contents

| Section | | Page |
|---|---|---|
| SI.1 | Model for the dynamics of the flow in an open-channel with multi trigger valves | S2 |
| SI.2 | Influence of the length of the side channel | S8 |
| SI.3 | Side channels located oppositely and facing each other | S9 |
| SI.4 | Engineering drawings of trigger valve devices | S12 |
| SI.5 | Additional experimental data | S14 |
| SI.6 | Supporting information references | S20 |

#### Additional supporting information included:

Video: flow experiment with trigger valves on the same side (.mp4)
Design File: trigger valves on the same side (.STL)
Design File: trigger valves on opposing sides (.STL)
Design File: trigger valves 15 mm apart (.STL)
Design File: trigger valves 30 mm apart (.STL)
Design File: trigger valves extended exit length (.STL)
Design File: trigger valves side view imaging (.STL)
Design File: triggervalves nitrite detection (.STL)
Code: main_SSv7_meniscus (.m)
Code: meniscus_big_loop_SSv7 (.m)
Code: main_SSv7_meniscus (.m)
Code: meniscus_big_loop_SSv8 (.m)
Code: main_SSv11_long (.m)
Data: raw data file_v2 (.xlsx)

### SI.1. Model for the dynamics of the flow in an open-channel with multi trigger valves

#### 1. Geometry

A sketch of the device we are considering here is shown in Figure SI.1.1. The valves can be placed on the same side of the channel or on both sides. The type of arrangement does not change the dynamics of the flow (in the case of miscible liquids), but it changes the arrangement of the stream tubes. In this SI, we only analyze the dynamic of the flow.

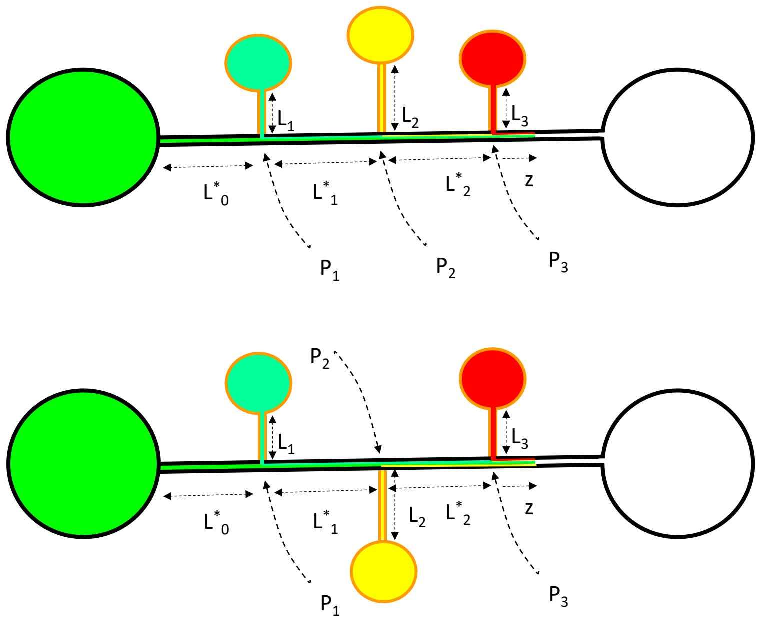

**Fig.SI.1.1**. Sketch of the main channel with the trigger valves.

#### 2. Notations

The notations used in this section are listed in the table 1.

**Table 1**: Notations

| Name | notation | unit | Remarks/References |
|---|---|---|---|
| Length | $L_i$ | mm | Lengths of side channels |
| Length | $L^*_i$ | mm | Lengths between two nodesin the main channel |
| Width | $w^*=w_0$ | mm | Width of main channel (uniform) |
| Width | $w_i$ | mm | Widths of side channels |
| Depth | $h^*=h_0$ | mm | Depth of main channel (uniform) |
| Depth | $h_i$ | mm | Depths of side channels |
| Cross-sectional area | $S^*=S_0$ | $mm^2$ | Main channel (uniform) |
| Cross-sectional area | $S_i$ | $mm^2$ | side channels |
| Average friction length | $\bar{\lambda}_\iota^*=\lambda_0$ | mm | Main channel (uniform)[1] |
| Average friction length | $\bar{\lambda}_\iota$ | mm | Side channel[1] |

| | | | |
|---|---|---|---|
| Perimeter (total) | $p^*=p_i$ | mm | Main channel (uniform) |
| Perimeter (total) | $p_i$ | mm | Side channel, Includes free perimeter |
| Perimeter (wetted) | $p_w^*=p_{W,0}$ | mm | Main channel, only wetted part |
| Perimeter (wetted) | $p_{W,i}$ | mm | Side channel, only wetted part |
| Viscosity (dynamic) | $\mu$ | mPa.s | [for nonanol |
| Surface tension | $\gamma$ | mN/m | for nonanol |
| Contact angle | $\theta$ | rad | Static contact angle |
| Generalized Cassie angle | $\theta^*_i$ | rad | Generalized Cassie angle for open channels |
| Contact angle | $\theta_d$ | rad | Dynamic contact angle |
| Pressure | $P$ | mPa | Pressure at the nodes |
| Capillary pressure | $P_{cap}$ | mPa | Laplace pressure of the meniscus |
| Velocity | $V_i$ | mm/s | Averaged in a cross section (side channels) |
| Velocity | $V_i^*$ | mm/s | Averaged in a cross section (main channel) |
| Travel distance | $z$ | mm | Middle of advancing meniscus |
| Coefficient of TCL friction | $\zeta$ | Pa.s | MKT approach [2] |
| Geometrical coefficient | $A_i$ | mm³ | Side channel |
| Geometrical coefficient | $A*$ | mm³ | Main channel |
| Auxiliary geometrical coefficient | $B_i$ | mm³ | Node coefficients |

#### 3. Inlet main channel

The first part of the flow occurs in the inlet channel, before the first trigger valve. In the case of nonanol—which presents a relatively high viscosity—, the duration of the inertial regime is very short, less than 10 ms[3–5], and the length of the channel affected by the initial inertial regime is less than 1 to 2 mm. So, we consider here only the viscous regime, described by Lucas, Washburn and Rideal (LWR law) in the years 1928-1930[2, 6–8] and recently generalized to arbitrary channels[1, 9–11]

$$z_0(t) = \sqrt{\frac{\gamma}{\mu} 2\bar{\lambda}_0 \cos\theta_0^*\ t}, \tag{SI1.1}$$

where t is the time ($t$=0, $z$=0), and $\theta_0^*$ is the so-called generalized Cassie angle[12] given by the relation

$$\cos\theta_0^* = \frac{p_W}{p}\cos\theta_0 - \frac{p_F}{p}, \tag{SI1.2}$$

where $p_W$ and $p_F$ are respectively the wetted and free perimeter in a cross section. For a rectangular channel, $p_W=2h+w$ and $p_F=w$. A correction may be done to the preceding relation at the channel inlet where the fluid velocity is still high (even in the viscous regime). A dynamic contact angle (DCA) should be taken into account. It has been shown that in open channel geometry the molecular kinetic theory (MKT) provides an accurate value for the DCA.[5, 13–16]

The correction for the DCA is

$$\cos\theta_{d,0} = \cos\theta_0 - \frac{\zeta}{\gamma} V \tag{SI1.3}$$

where $\zeta$ is the coefficient of triple contact line (TPL) friction. This expression modifies the generalized Cassie angle

$$\cos\theta_{d,0}^* = \frac{p_W}{p}\cos\theta_{d,0} - \frac{p_F}{p} = \left(\frac{p_W}{p}\cos\theta_0 - \frac{p_F}{p}\right) - \frac{p_W}{p}\frac{\zeta}{\gamma} V = \cos\theta_0^* - \frac{p_W}{p}\frac{\zeta}{\gamma} V \tag{SI1.4}$$

The generalized LWR law is then[4]

$$z = -\bar{\lambda}_0 \frac{p_{w,0}}{p_0}\frac{\zeta}{\mu} + \sqrt{\left(\bar{\lambda}_0 \frac{p_{w,0}}{p_0}\frac{\zeta}{\mu}\right)^2 + z_0^2(t)}\,. \tag{SI1.5}$$

The front meniscus reaches the first TGV at the time $t_0$ determined by

$$t_0 = \frac{\left(L_0^* + \bar{\lambda}_0 \frac{p_{w,0}}{p_0}\frac{\zeta}{\mu}\right)^2 - \left(\bar{\lambda}_0 \frac{p_{w,0}}{p_0}\frac{\zeta}{\mu}\right)^2}{\frac{\gamma}{\mu} 2\bar{\lambda}_0 \cos\theta_{d,0}^*} . \quad \text{(SI1.6)}$$

#### 4. First trigger valve

A sketch of the flow after the first trigger valve (and before the second TGV) is shown in Figure SI.1.2. It is assumed here that the first TGV is far enough from inlet reservoir to neglect the dynamic contact angle. There is a very short inertial burst when a TGV opens, but it is evanescent and we don't take it into account here.

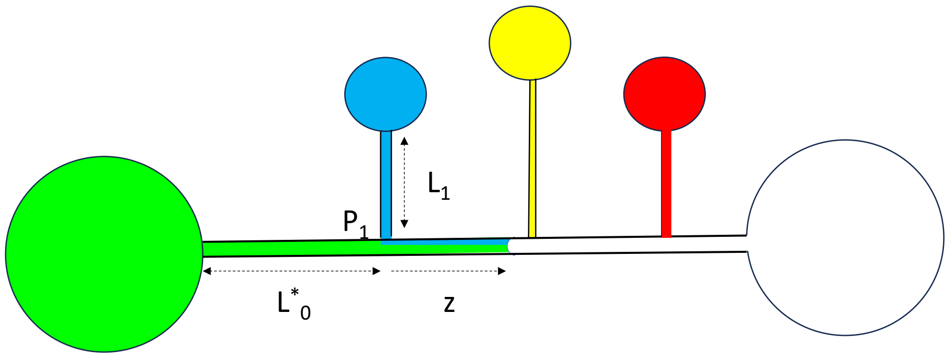

**Figure SI.1.2**. Sketch of the device with the advancing meniscus between the first and second TG. The "*" characterizes the main channel, while the side channels are noted *i*=1,…n.

The pressure $P_1$ may be expressed by the two relations [4]

$$P_1 = \frac{L_1\, p_1}{S_1}\, \mu \frac{V_1}{\bar{\lambda}_1} = \mu \frac{S_1 V_1}{A_1}, \quad \text{(SI1.7)}$$

considering the side channel #1, where $A_1 = \frac{\bar{\lambda}_1 S_1^2}{L_1\, p_1}$, and

$$P_1 = \frac{L_0^*\, p_0}{S_0}\, \mu \frac{V_0^*}{\bar{\lambda}_0} = \mu \frac{S_0 V_0^*}{A_0^*}, \quad \text{(SI1.8)}$$

considering the main channel (index 0), where $A_0^* = \frac{\bar{\lambda}_0 S_0^2}{L_0^*\, p_0}$. The mass conservation equation yields the relation

$$S_0 V_1^* = S_0 V_0^* + S_1 V_1. \quad \text{(SI1.9)}$$

Multiplying (SI1.7) by $S_1$ and (SI1.8) by $S_0$, adding the result and using (SI1.9) leads to

$$S_0 V_1^* = S_0 V_0^* + S_1 V_1 = \frac{P_1}{\mu}\left(A_0^* + A_1\right). \quad \text{(SI1.10)}$$

Let us denote $B_0 = A_0^* = \frac{\bar{\lambda}_0 S_0^2}{L_0^*\, p_0}$. The pressure $P_1$ is then given by

$$P_1 = \mu \frac{S_0 V_1^*}{(B_0+A_1)} = \mu \frac{Q_1^*}{(B_0+A_1)} \ , \tag{SI1.11}$$

where $Q_1^*$ is the volumetric flow rate in the main channel after the first TGV. In the viscous regime, the equation for the fluid motion is given by the balance between capillary pressure and pressure drop. If *z* denotes the distance taken from the first node (first TGV), we have

$$P_{cap} = \frac{\gamma\, p_0 \cos\theta_0^*}{S_0} = \frac{z\, p_0}{S_0} \mu \frac{V_1^*}{\bar{\lambda}_0} + P_1 = \frac{z\, p_0}{S_0} \mu \frac{V_1^*}{\bar{\lambda}_0} + \mu \frac{S_0 V_1^*}{(B_0+A_1)} = \mu \frac{z}{L_0^*} \frac{S_0 V_1^*}{A_0^*} + \mu \frac{S_0 V_1^*}{(B_0+A_1)}.\ . \tag{SI1.12}$$

Inserting $V_1^* = dz/dt$ in (SI1.12) produces the differential equation

$$\frac{1}{2} \frac{1}{L_0^* A_0^*} \frac{dz^2}{dt} + \frac{1}{(B_0+A_1)} \frac{dz}{dt} - \frac{\gamma}{\mu} \frac{p_0}{S_0^2} \cos\theta_0^* = 0 \ , \tag{SI1.13}$$

or,

$$\frac{dz^2}{dt} + 2 \frac{L_0^* A_0^*}{(B_0+A_1)} \frac{dz}{dt} - \frac{\gamma}{\mu} 2\bar{\lambda}_0 \cos\theta_0^* = 0 \ . \tag{SI1.14}$$

Remarking that the travel distance in the main channel without TGVs is $z_0 = \sqrt{\frac{\gamma}{\mu} 2\bar{\lambda}_0 \cos\theta_0^*\, \tau_1}$, where $\tau_1$ is the time taken after the passage of the first TGV ($\tau_1 = t - t_0$), integration of (SI1.14) leads to the solution

$$z = -\frac{L_0^* A_0^*}{(B_0+A_1)} + \sqrt{\left[\frac{L_0^* A_0^*}{(B_0+A_1)}\right]^2 + z_0^2} \ . \tag{SI1.15}$$

At the time $t_1$, the front meniscus reaches the second TGV. This time is given by

$$t_1 = \frac{\left(L_1^* + \frac{L_0^* A_0^*}{(B_0+A_1)}\right)^2 - \left(\frac{L_0^* A_0^*}{(B_0+A_1)}\right)^2}{\frac{\gamma}{\mu} 2\bar{\lambda}_0 \cos\theta_0^*} + t_0 = \frac{L_1^{*2} + \frac{2 L_1^* L_0^* A_0^*}{(B_0+A_1)}}{\frac{\gamma}{\mu} 2\bar{\lambda}_0 \cos\theta_0^*} + t_0. \tag{SI1.16}$$

In the case of the main channel alone—without any TGV— the geometrical coefficient $A_1$=0, and relation (SI1.16) reduces to $t_1 = \frac{(L_1^* + L_0^*)^2}{\frac{\gamma}{\mu} 2\bar{\lambda}_0 \cos\theta_0^*}$, which is the generalized LWR expression.

#### 5. Second trigger valve

Let us now express the pressure $P_2$. On one hand,

$$P_2 = \frac{L_2\, p_2}{S_2} \mu \frac{V_2}{\bar{\lambda}_2} = \mu \frac{S_2 V_2}{A_2} = \mu \frac{Q_2}{A_2}, \tag{SI1.17}$$

considering the side channel #2. Considering the main channel

$$P_2 = P_1 + \frac{L^*_1 p_0}{S_0} \mu \frac{V_1^*}{\bar{\lambda}_0} = P_1 + \mu \frac{S_0 V_1^*}{A_1^*} = P_1 + \mu \frac{Q_1^*}{A_1^*}, \tag{SI1.18}$$

where $A_1^* = \frac{\bar{\lambda}_0 S_0^2}{L^*_1 p_0}$. Substituting (SI1.11) in (SI1.18) yields

$$P_2 = \mu \frac{S_0 V_1^*}{(B_0+A_1)} + \mu \frac{S_0 V_1^*}{A_1^*} = \mu S_0 V_1^* \left[\frac{1}{(B_0+A_1)} + \frac{1}{A_1^*}\right] = \mu \frac{S_0 V_1^*}{B_1} \ , \tag{SI1.19}$$

where $B_1 = \frac{1}{\left[\frac{1}{(B_0+A_1)}+\frac{1}{A_1^*}\right]}$. Remembering that the mass conservation equation at node 2 is

$$S_0 V_2^* = S_0 V_1^* + S_2 V_2 = S_0 V_0^* + S_1 V_1 + S_2 V_2 \,, \qquad \text{(SI1.20)}$$

and adding (SI1.17) and (SI1.19)

$$S_0 V_2^* = S_0 V_1^* + S_2 V_2 = \frac{P_2}{\mu} B_1 + \frac{P_2}{\mu} A_2 = \frac{P_2}{\mu}(A_2 + B_1). \qquad \text{(SI1.21)}$$

Again, the equation for the fluid motion is given by the balance between capillary pressure and pressure drop

$$P_{cap} = \frac{\gamma\, p_0 \cos\theta_0^*}{S_0} = \frac{z\, p_0}{S_0}\mu \frac{V_2^*}{\bar{\lambda}_0} + P_2 = \frac{z}{L_0^*}\mu \frac{S_0 V_2^*}{A_0^*} + \mu \frac{S_0 V_2^*}{(A_2+B_1)}. \qquad \text{(SI1. 22)}$$

Remembering that $V_2^* = dz/dt$, we obtain the differential equation

$$\frac{1}{2\, L_0^* A_0^*}\frac{dz^2}{dt} + \frac{1}{(A_2+B_1)}\frac{dz}{dt} - \frac{\gamma}{\mu}\frac{p_0}{S_0^2}\cos\theta_0^* = 0 \;, \qquad \text{(SI1.23)}$$

Still using the travel distance in the main channel without TGV; $z_0 = \sqrt{\frac{\gamma}{\mu} 2\bar{\lambda}_0 \cos\theta_0^* \tau_2}$, where $\tau_2$ is the time taken after the passage of the second TGV ($\tau_2 = t - t_1$), integration of (SI1.23) leads to the solution

$$z = -\frac{L_0^* A_0^*}{(A_2+B_1)} + \sqrt{\left[\frac{L_0^* A_0^*}{(A_2+B_1)}\right]^2 + z_0^2}\,. \qquad \text{(SI1.24)}$$

The same approach as precedingly produces the time $t_2$ when the meniscus reaches the third TGV

$$t_2 = \frac{\left(L_2^* + \frac{L_0^* A_0^*}{(A_2+B_1)}\right)^2 - \left(\frac{L_0^* A_0^*}{A_2+B_1}\right)^2}{\frac{\gamma}{\mu} 2\bar{\lambda}_0 \cos\theta_0^*} + t_1 = \frac{L_2^{*2} + \frac{2 L_2^* L_0^* A_0^*}{(A_2+B_1)}}{\frac{\gamma}{\mu} 2\bar{\lambda}_0 \cos\theta_0^*} + t_1 \;. \qquad \text{(SI1.25)}$$

#### 6. Third trigger valve

The pressure $P_3$ is obtained similarly

$$P_3 = \frac{L_3\, p_3}{S_3}\mu \frac{V_3}{\bar{\lambda}_3} = \mu \frac{S_3 V_3}{A_3} = \mu \frac{Q_3}{A_3}, \qquad \text{(SI1.26)}$$

considering the side channel #3, and considering the main channel

$$P_3 = P_2 + \frac{L_2^*\, p_0}{S_0}\mu \frac{V_2^*}{\bar{\lambda}_0} = P_2 + \mu \frac{S_0 V_2^*}{A_2^*}, \qquad \text{(SI1.27)}$$

where $A_2^* = \frac{\bar{\lambda}_0 S_0^2}{L_2^*\, p_0}$. Substituting (SI1.21) in (SI1.27) yields

$$P_3 = \mu \frac{S_0 V_2^*}{(A_2+B_1)} + \mu \frac{S_0 V_2^*}{A_2^*} = \mu\, S_0 V_2^* \left\{\frac{1}{(A_2+B_1)} + \frac{1}{A_2^*}\right\} = \frac{\mu\, S_0 V_2^*}{B_2} \;, \qquad \text{(SI1.28)}$$

where $B_2 = \frac{1}{\left[\frac{1}{(B_1+A_2)}+\frac{1}{A_2^*}\right]}$. Remembering that the mass conservation equation at node 3 is

$$S_0V_3^* = S_0V_2^* + S_3V_3 = S_0V_0^* + \sum_{i=1,3} S_iV_i \;, \qquad \text{(SI1.29)}$$

and adding (SI1.26) and (SI1.28)

$$S_0V_3^* = S_0V_2^* + S_3V_3 = \frac{P_3}{\mu} B_2 + \frac{P_3}{\mu} A_3 = \frac{P_3}{\mu} (A_3 + B_2), \qquad \text{(SI1.30)}$$

Finally, considering the part of the main channel where the front meniscus is advancing, the capillary pressure balances the pressure drop

$$P_{cap} = \frac{\gamma\, p_0 \cos\theta_0^*}{S_0} = \frac{z\, p_0}{S_0}\mu \frac{V_3^*}{\bar{\lambda}_0} + P_3 = \mu \frac{z}{L_0^*} \frac{S_0V_3^*}{A_0^*} + \mu \frac{S_0V_3^*}{(A_3+B_2)}. \qquad \text{(SI1.31)}$$

Substituting $V_3^* = dz/dt$ in (SI1.31) yields the differential equation

$$\frac{1}{2L_0^*A_0^*} \frac{dz^2}{dt} + \frac{1}{(A_3+B_2)} \frac{dz}{dt} - \frac{\gamma}{\mu} \frac{p_0}{S_0^2} \cos\theta_0^* = 0 \;, \qquad \text{(SI1.32)}$$

Still using the travel distance in the main channel without TGVs $z_0 = \sqrt{\frac{\gamma}{\mu} 2\bar{\lambda}_0 \cos\theta_0^* \tau_3}$, where $\tau_3$ is the time taken after the passage of the third TG ($\tau_3 = t - t_2$), integration of (SI1.32) leads to the solution

$$z = -\frac{L_0^*A_0^*}{(A_3+B_2)} + \sqrt{\left[\frac{L_0^*A_0^*}{A_3+B_2}\right]^2 + z_0^2} \;. \qquad \text{(SI1.33)}$$

#### 7. Generalization

The travel distance of the front meniscus is given by the recurrence relation

$$z = -\frac{L_0^*A_0^*}{(A_n+B_{n-1})} + \sqrt{\left[\frac{L_0^*A_0^*}{A_n+B_{n-1}}\right]^2 + \frac{\gamma}{\mu} 2\bar{\lambda}_0 \cos\theta_0^* (t - t_{n-1})} \;, \qquad \text{(SI1.34)}$$

where $A_n = \frac{\bar{\lambda}_n S_n^2}{L_n p_n}$ and $B_0 = A_0^* = \frac{\bar{\lambda}_0 S_0^2}{L_0^*\, p_0}$, $B_1 = \frac{1}{\left[\frac{1}{(B0+A_1)} + \frac{1}{A_1^*}\right]}$, and for all $n$: $B_n = \frac{1}{\left[\frac{1}{(B_{n-1}+A_n)} + \frac{1}{A_n^*}\right]}$. On the other hand, the times when the meniscus reaches a TGV is

$$t_n = \frac{L_n^{*2} + \frac{2L_n^*L_0^*A_0^*}{p_0(A_n+B_{n-1})}}{\frac{\gamma}{\mu} 2\bar{\lambda}_0 \cos\theta_0^*} + t_{n-1} \;. \qquad \text{(SI1.35)}$$

Finally, the general expression of the velocity of the front meniscus is

$$V = \frac{1}{2} \frac{\frac{\gamma}{\mu} 2\bar{\lambda}_0 \cos\theta_0^*}{\sqrt{\left[\frac{L_0^*A_0^*}{B_{n-1}+A_n}\right]^2 + \frac{\gamma}{\mu} 2\bar{\lambda}_0 \cos\theta_0^* (t - t_{n-1})}} = \tilde{V}_0 \frac{\tilde{z}_0}{z + \frac{L_0^*A_0^*}{(A_n+B_{n-1})}}, \qquad \text{(SI1.36)}$$

where $\tilde{z}_0 = \sqrt{\frac{\gamma}{\mu} 2\bar{\lambda}_0\, cos\;\theta_0^*\; t}$ and $\tilde{V}_0 = \frac{\frac{\gamma}{\mu} 2\bar{\lambda}_0 \cos\theta_0^*}{\tilde{z}_0}$, corresponding to the main channel without TGVs.

#### SI.2. Influence of the length of the side channel

In this section, we investigate the effect of the side channel length on the flow after the TGV. The liquid is nonanol and the characteristic dimensions are listed in table SI2

Table SI2

| Inlet channel | Side channel #1 | Side channel #2 | Side channel #3 |
|---|---|---|---|
| $L_0$= 80 mm | $L_1$ = 2 mm | $L_1$ = 8 mm | $L_1$ = 80 mm |
| $w_0$ = 1mm | $w_1$ = 0.4 mm | $w_1$ = 0.4 mm | $w_1$ = 0.4 mm |
| $h_0$=1mm | $h_1$ = 0.6 mm | $h_1$ = 0.6 mm | $h_1$ = 0.6 mm |

The results obtained using the model are shown in Figure SI.2.1. A short side channel ($L_1 << L_0$) produces a jump of the velocity at the TG, as shown by the green line in figure SI.2. This jump decreases when the length of the side channel increases. When the two upstream lengths are equal ($L_1 = L_0$), the jump disappears, as shown by the purple line in Figure SI.2.1.

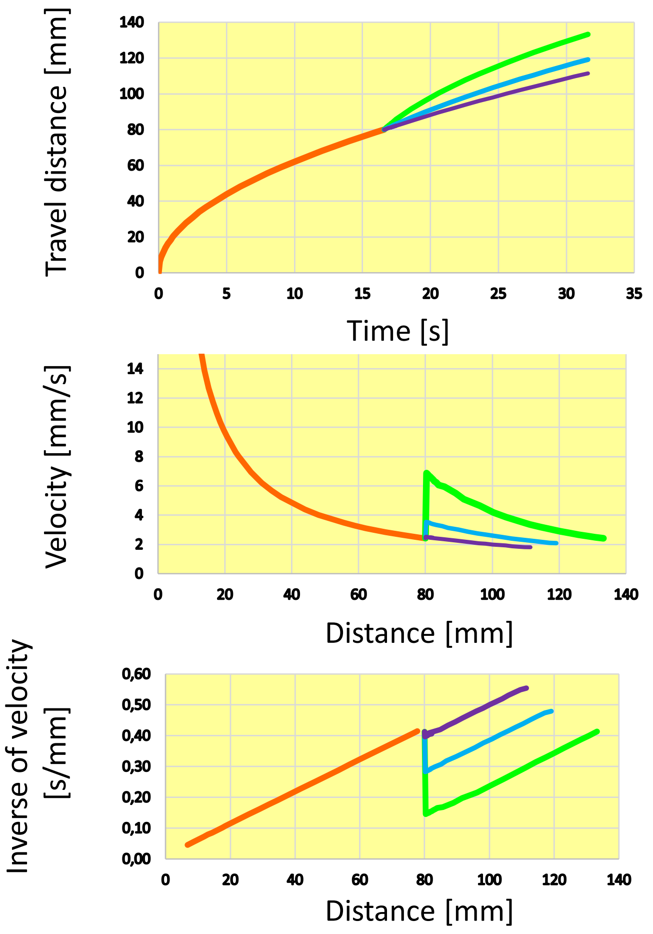

**Figure SI.2.1.** Top, travel distance vs time; middle, velocity vs distance in channel; bottom, inverse of velocity vs distance in channel. The orange line corresponds to the inlet channel, the green line to the side channel #1, the blue line to side channel #2, the purple line to side channel #3.

#### SI.3. Side channels located opposite and facing each other

In this section, we write the model for the dynamics (travel distance vs. time) of the flow in the case of oppositely facing side channels (Figure SI.3.1).

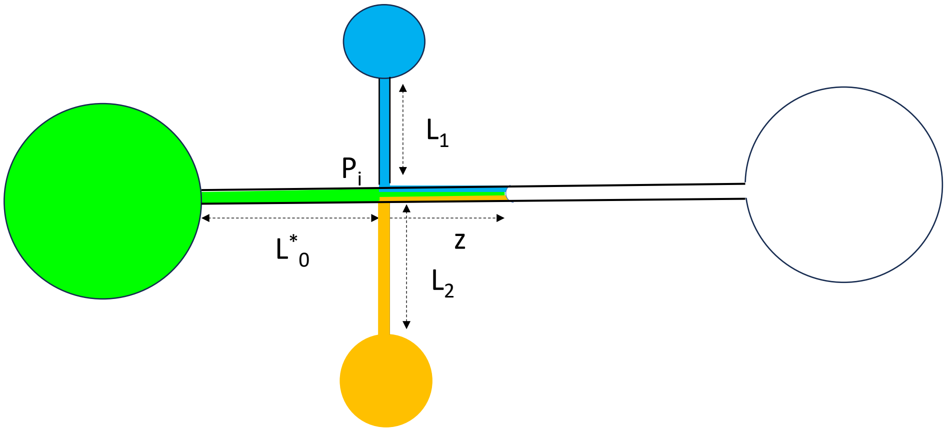

**Figure SI.3.1.** Sketch of the trigger-valve channel with opposite TGs.

##### 1. Model

The pressure at the intersection of the three channels is denoted $P_i$, which is the pressure of the node at the intersection of the three channels. Let us first write the pressure drop in the three channels ending at the intersection

$$P_i = \frac{p_0 L_0^*}{S_0}\,\mu \frac{V_0}{\bar{\lambda}_0}\ , \qquad \text{(SI.3.1)}$$

for the main channel (index *0*), and

$$P_i = \frac{p_1\, L_1}{S_1}\,\mu \frac{V_1}{\bar{\lambda}_1}\ , \qquad \text{(SI.3.2)}$$

for the first side channel (index *1*), and

$$P_i = \frac{p_2\, L_2}{S_2}\,\mu \frac{V_2}{\bar{\lambda}_2}\ , \qquad \text{(SI.3.3)}$$

for the second side channel (index 2). The mass conservation equation yields

$$S_0 V = S_0 V_0 + S_1 V_1 + S_2 V_2\ . \qquad \text{(SI.3.4)}$$

In the viscous regime, the balance between capillary force and wall friction is [6,17]

$$\frac{p_0\,\gamma\, cos\theta_0^*}{S_0} = \frac{p_0\, z}{S_0}\,\mu \frac{V}{\bar{\lambda}_0} + P_i\ , \qquad \text{(SI.3.5)}$$

where $cos\theta_0^*$ is the cosine of the generalized Cassie angle. Finally, we consider the additional relation

$$V = \frac{dz}{dt} \quad . \tag{SI.3.6}$$

Hence, we have 6 equations (SI.3.1 to SI.3.6) for the 6 unknowns $\{V, V_0, V_1, V_2, z, P_i\}$. Multiplying (SI.3.1) by $S_0$, (SI.3.2) by $S_1$, (SI.3.3) by $S_2$, and substituting the result in (SI.3.4) yields

$$S_0 V = \frac{P_i}{\mu}\left(\frac{\bar{\lambda}_0 S_0}{p_0\, L_0} + \frac{\bar{\lambda}_1 S_1}{p_1\, L_1} + \frac{\bar{\lambda}_2 S_2}{p_2\, L_2}\right) = \frac{P_i}{\mu}(A_0 + A_1 + A_2) \quad . \tag{SI.3.7}$$

Substituting (SI.3.6) and (SI.3.7) in SI.3.5) yields

$$\frac{p_0\, \gamma\, cos\theta_0^*}{S_0} = \frac{p_0}{2S_0}\, \mu \frac{1}{\bar{\lambda}_0} \frac{dz^2}{dt} + \mu \frac{S_0}{(A_0+A_1+A_2)} \frac{dz}{dt} \quad . \tag{SI.3.8}$$

Time integration of (SI.3.8) leads to a quadratic expression for $z$

$$z^2 + \left(\frac{2\bar{\lambda}_0 S_0^{\,2}}{p_0\, \sum_{i=0,2} A_i}\right) z - \frac{\gamma\, 2\bar{\lambda}_0\, cos\theta_0^*}{\mu}\tau = 0 \ , \tag{SI.3.9}$$

where $\tau$ is the time counted from the passage of the meniscus at the intersection. The expression for the travel distance $z$ vs. time $\tau$ is then

$$z = -\left(\frac{\bar{\lambda}_0 S_0^{\,2}}{p_0\, \sum_{i=0,2} A_i}\right) + \sqrt{\left(\frac{\bar{\lambda}_0 S_0^{\,2}}{p_0\, \sum_{i=0,2} A_i}\right)^2 + \frac{\gamma\, 2\bar{\lambda}_0\, cos\theta_0^*}{\mu}\tau} \quad . \tag{SI.3.10}$$

In consequence, the velocity of the meniscus is

$$V = \frac{\frac{\gamma\, \bar{\lambda}_0\, cos\theta_0^*}{\mu}}{\sqrt{\left(\frac{\bar{\lambda}_0 S_0^{\,2}}{p_0\, \sum_{i=0,2} A_i}\right)^2 + \frac{\gamma\, 2\bar{\lambda}_0\, cos\theta_0^*}{\mu}\tau}} = \frac{\frac{\gamma\, \bar{\lambda}_0\, cos\theta_0^*}{\mu}}{z + \left(\frac{\bar{\lambda}_0 S_0^{\,2}}{p_0\, \sum_{i=0,2} A_i}\right)} \, . \tag{SI.3.11}$$

Remark that, if we note $z_0$ the travel distance in the main channel without side channels ($z_0 = \sqrt{\frac{\gamma\, 2\bar{\lambda}_0\, cos\theta_0^*}{\mu}\tau}$), and $\tilde{V}_0 = \frac{dz_0}{dt} = \frac{\gamma\, \bar{\lambda}_0\, cos\theta_0^*}{\mu}\frac{1}{z_0}$) we can rewrite (SI.3.11) as

$$\frac{V}{\tilde{V}_0} = \frac{z_0}{z + \left(\frac{\bar{\lambda}_0 S_0^{\,2}}{p_0\, \sum_{i=0,2} A_i}\right)} \, . \tag{SI.3.12}$$

#### 2. Example

Let us consider the same liquid (nonanol) and the dimensions listed in Table SI.3.1.

**Table SI.3.1**

| Inlet channel (index 0) | Side channel #1 | Side channel #2 |
|---|---|---|
| $L_0$= 80 mm | $L_1$ = 2 mm | $L_1$ = 2 mm |
| $w_0$ = 1mm | $w_1$ = 0.4 mm | $w_1$ = 0.4 mm |
| $h_0$=1mm | $h_1$ = 0.6 mm | $h_1$ = 0.6 mm |

The results obtained using the model (relations SI.3.10 and SI.3.11) are shown in Figure SI.3.2.

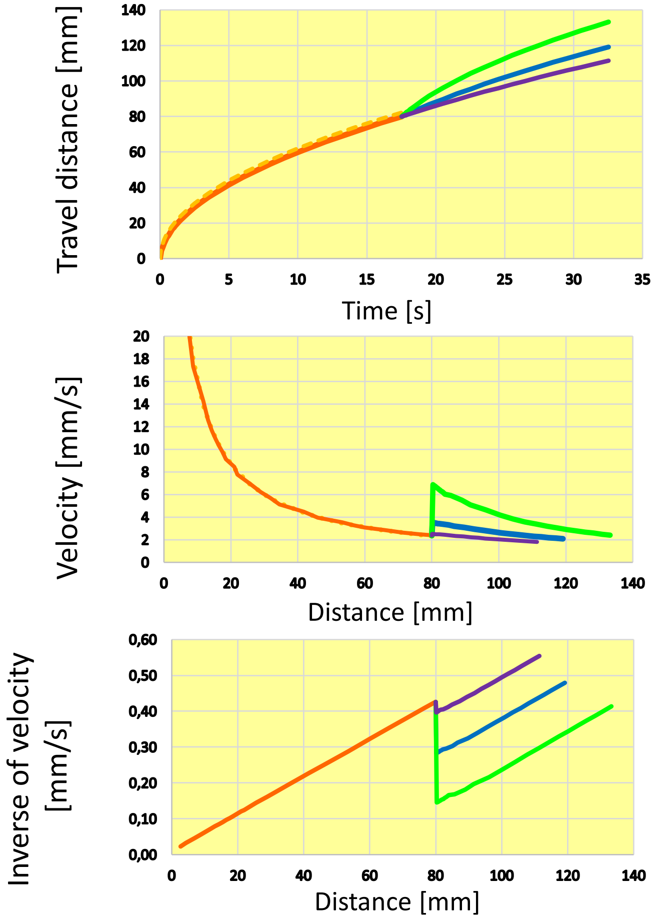

**Figure SI.3.2.** Top, travel distance vs time; middle, velocity vs distance in channel; bottom, inverse of velocity vs distance in channel. The orange line corresponds to the inlet channel, the green line to side channels of length 2 mm, the blue line to side channel of length 8 mm, and the purple line to side channels of length 80 mm.

#### SI.4. Engineering drawings of trigger valve devices

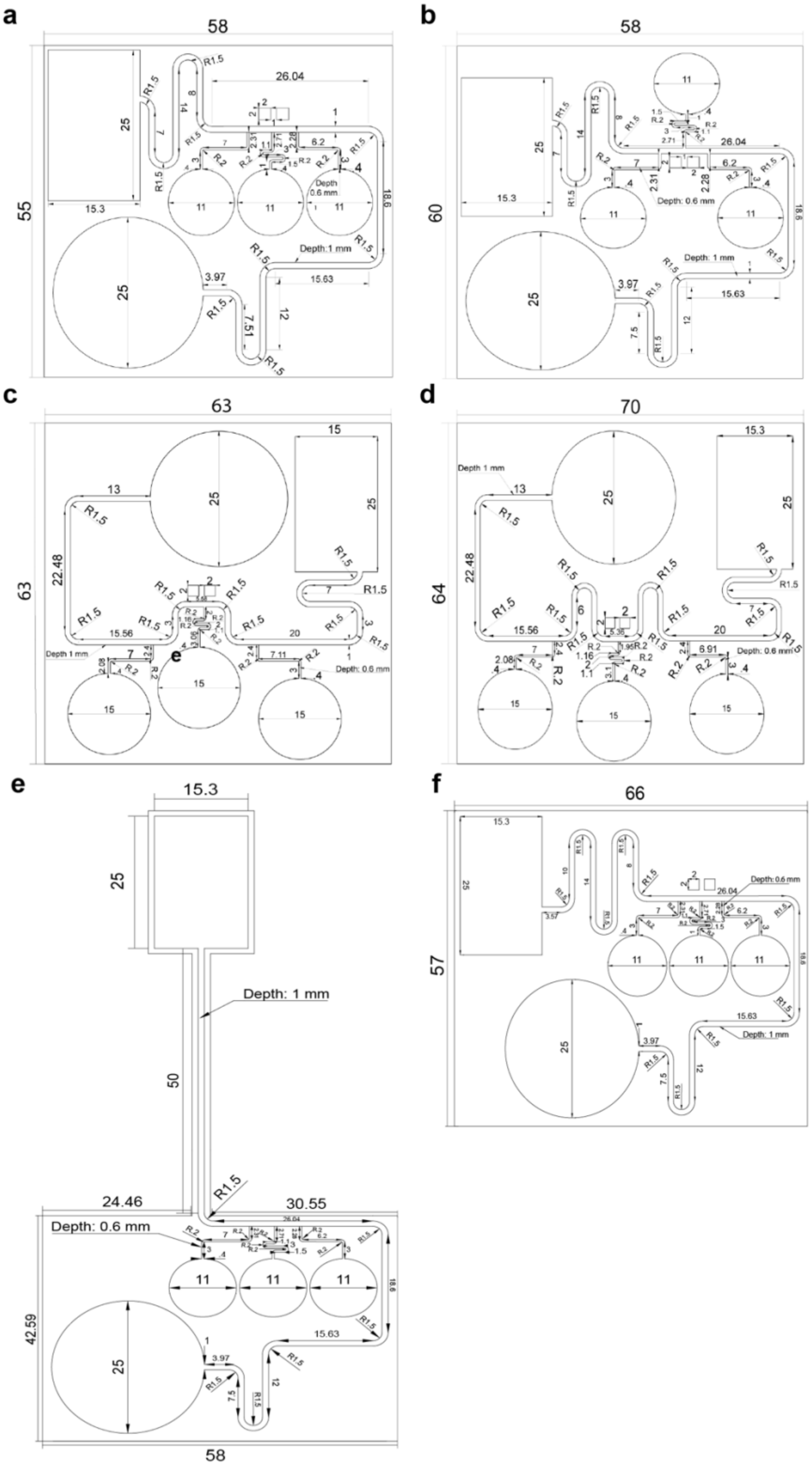

**Figure SI.4.1.** Engineering drawings of devices with channels on the same (a) and opposite sides (b) of the main channel and devices with 15 mm (c) and 30 mm (d) between trigger valves. Units in mm.

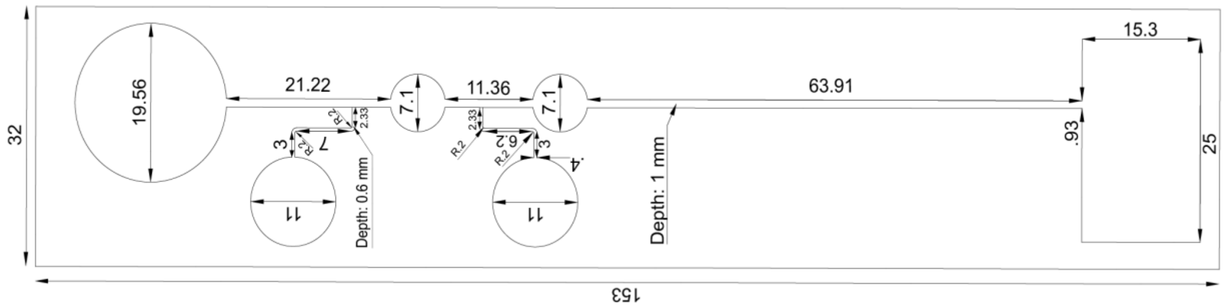

**Figure SI.4.2.** Engineering drawing of nitrite testing device. Units in mm.

#### SI.5. Additional experimental data

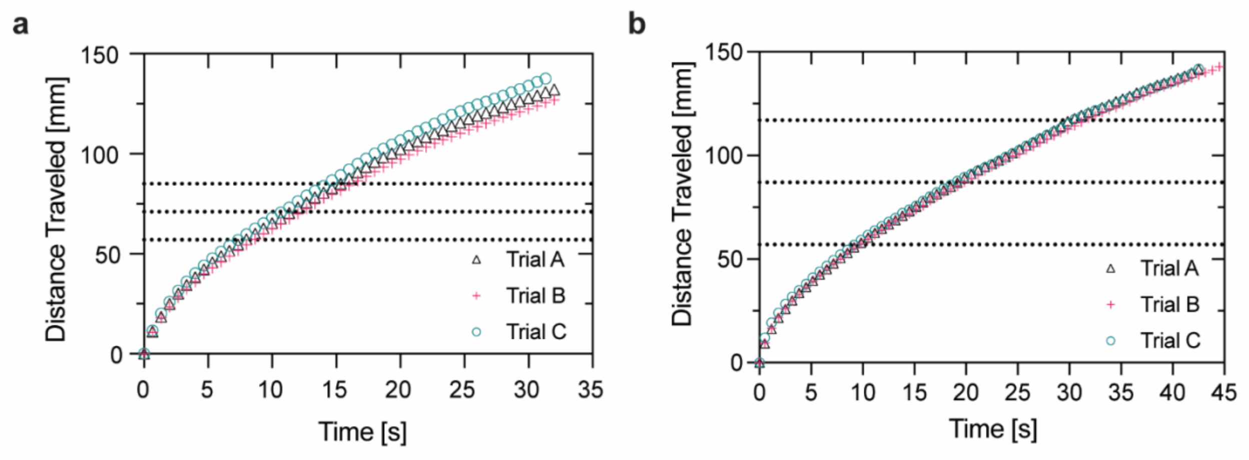

**Figure SI.5.1.** Travel distance data for all trials for programmed TGV release with valves separated by approximately 15 mm (a) and 30 mm (b). Every third data point is plotted for three trials (n=3).

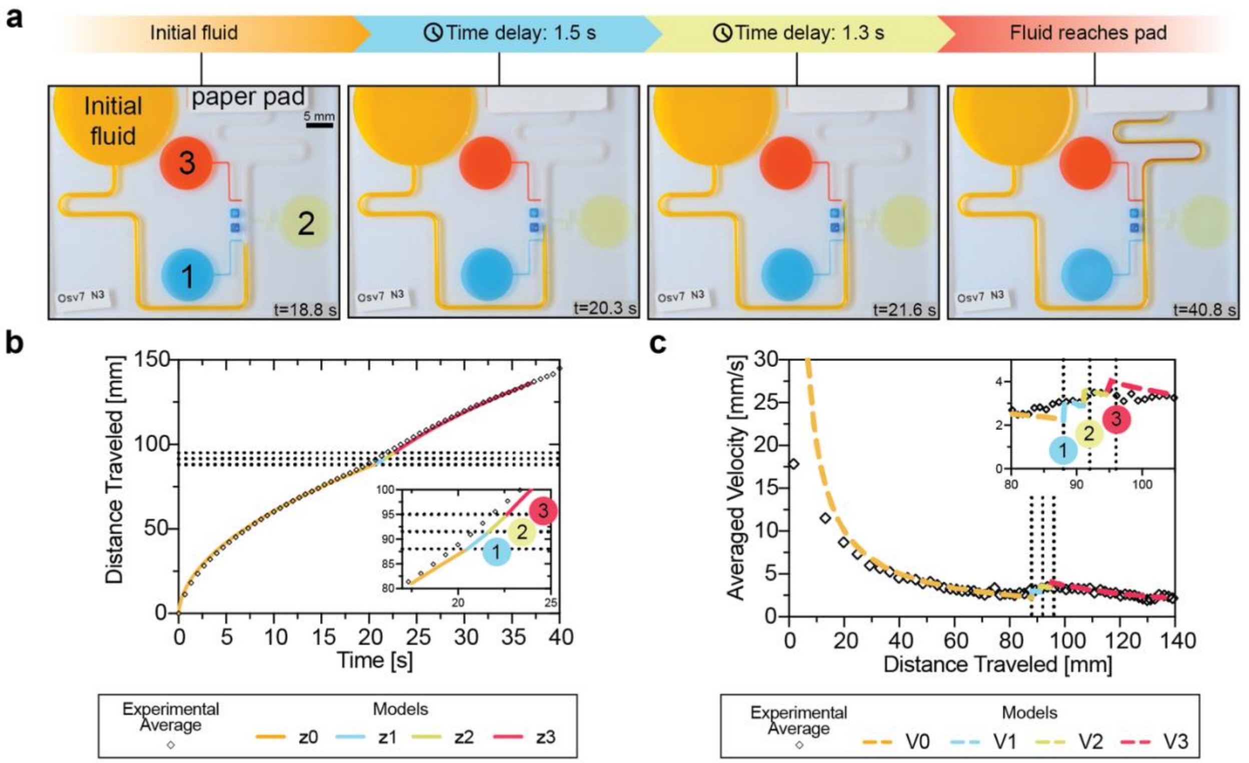

**Figure SI.5.2.** *Open trigger valves can be placed on opposite sides of main channel and maintain stair-like velocity trend.* Progression of fluid flow through device with shorter distances between oppositely-facing trigger valves (a). Comparison of the theoretical model (solid line) with experimental fluid front travel distance at the meniscus (black dots) for devices with three side channels on both sides of the main channel. Experimental data was averaged for three trials (n=3). Model is presented in segments corresponding to the calculated travel distances prior to the trigger valve release (orange, z0), between the first and second trigger valve (blue, z1), between the second the third valves (yellow, z2) and between the third valve and the paper pad (red, z3). Fluid velocities were calculated from the travel distance and the experimental data (diamonds) was compared against the calculated model (dashed line) velocities using a dynamic contact angle model (orange, V0) for the inlet. Model is shown at the first (blue, V1), second (yellow, V2), and third (red, V3) valves due to the increase in velocity upon trigger valve release.

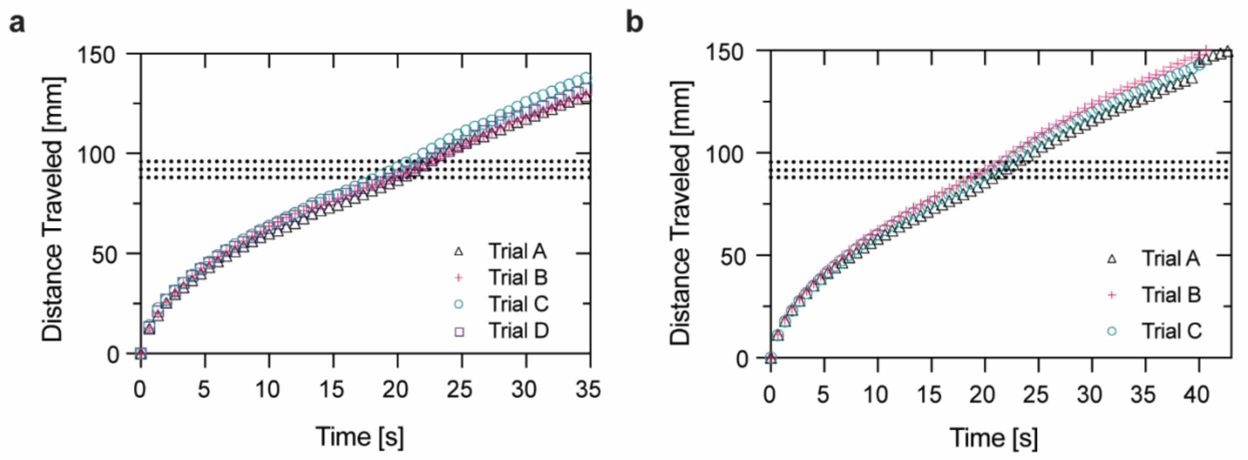

**Figure SI.5.3.** Travel distance data for all trials for TGV with shortened distances between valves on the same (a) and opposite sides (b) of the main channel. Every third data point is plotted for four (n=4) and three (n=3) trials for the TGVs on the same and opposite sides of the main channel, respectively.

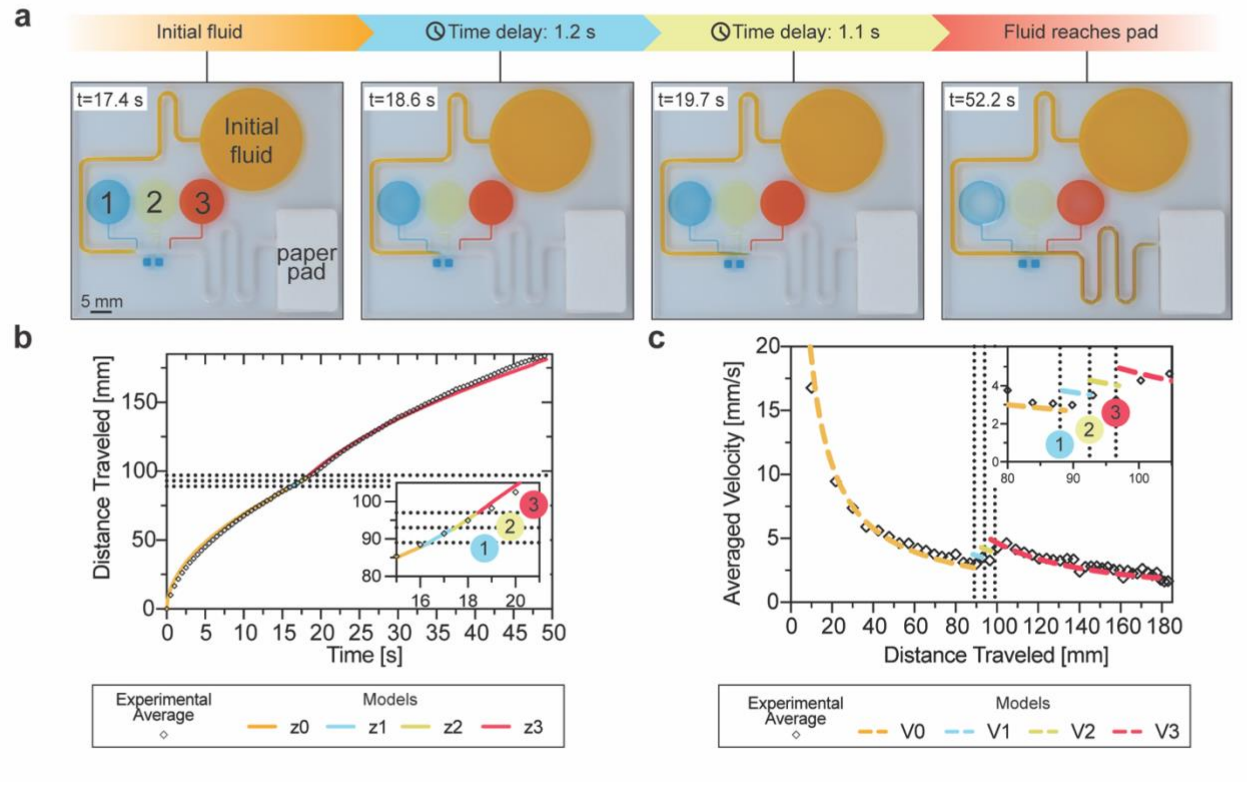

**Figure SI.5.4.** *Model predicts fluid velocity at approximately 80 mm after the last trigger valve.* Progression of fluid flow through device with a longer distance after the third trigger valve (a). Comparison of the theoretical model (solid line) with experimental fluid front travel distance at the meniscus (black dots) for devices with three side channels on the same side of the main channel with an extended channel length after the third valve. Experimental data was averaged for three trials (n=3). Model is presented in segments corresponding to the calculated travel distances prior to the trigger valve release (orange, z0), between the first and second trigger valve (blue, z1), between the second the third valves (yellow, z2) and between the third valve and the paper pad (red, z3). Fluid velocities were calculated from the travel distance and the experimental data (diamonds) was compared against the calculated model (dashed line) velocities using a dynamic contact angle model (orange, V0) for the inlet. Model is shown at the first (blue, V1), second (yellow, V2), and third (red, V3) valves due to the increase in velocity upon trigger valve release. Raw data can be found in Figure SI.5.5.

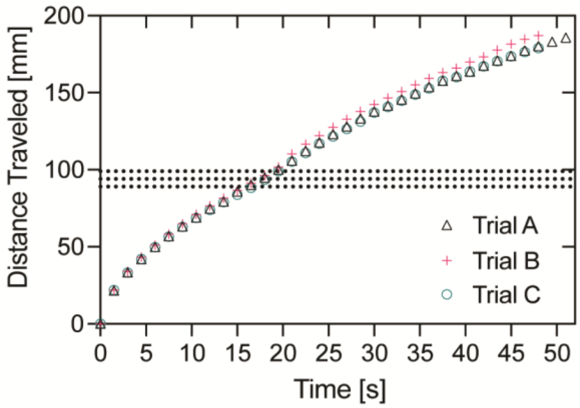

**Figure SI.5.5.** Travel distance data for TGV device with extended distance between last TGV and paper pad. Every third data point is plotted for three (n=3) trials.

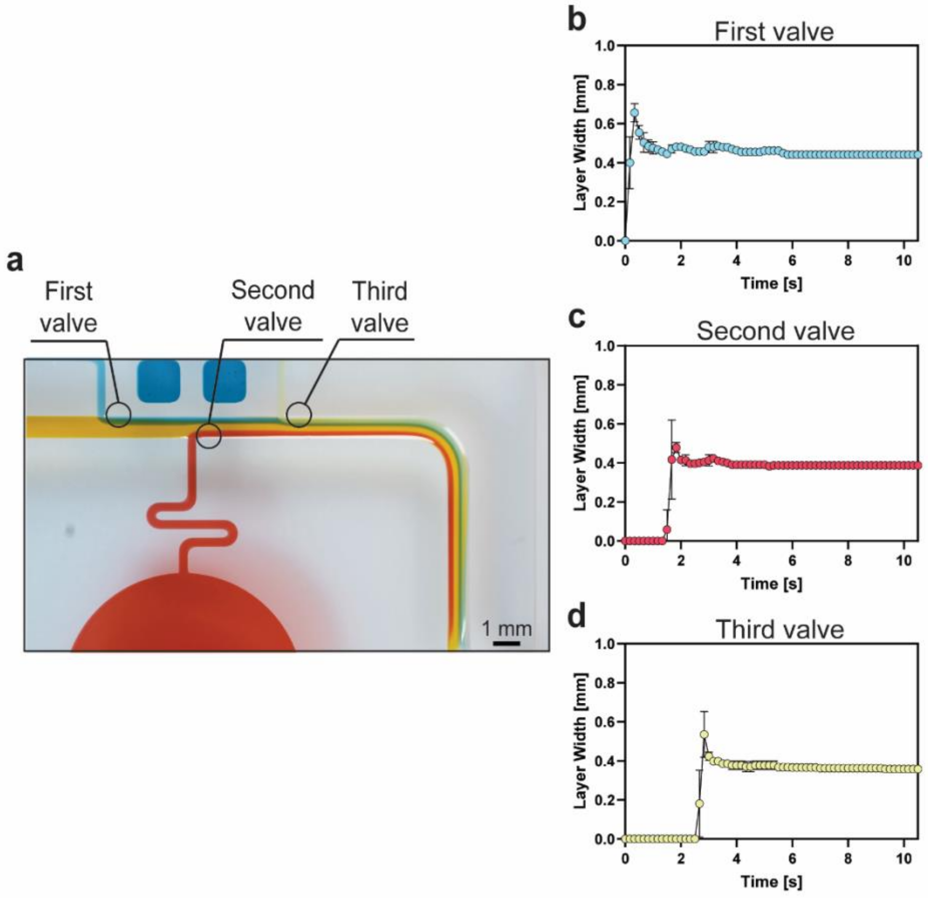

**Figure SI.5.6.** *Layers of trigger valves in oppositely facing configuration stabilize over time.* Image of the fluid layers in oppositely facing configuration (a). Circles indicate region where layer thickness measurements were taken for each layer. Plots of the measured layer widths over time for the parallel channel configuration after the first (c), second (d), and third (e) valves. Data is shown as an average of three replicates (n = 3) with error bars indicating standard deviation.

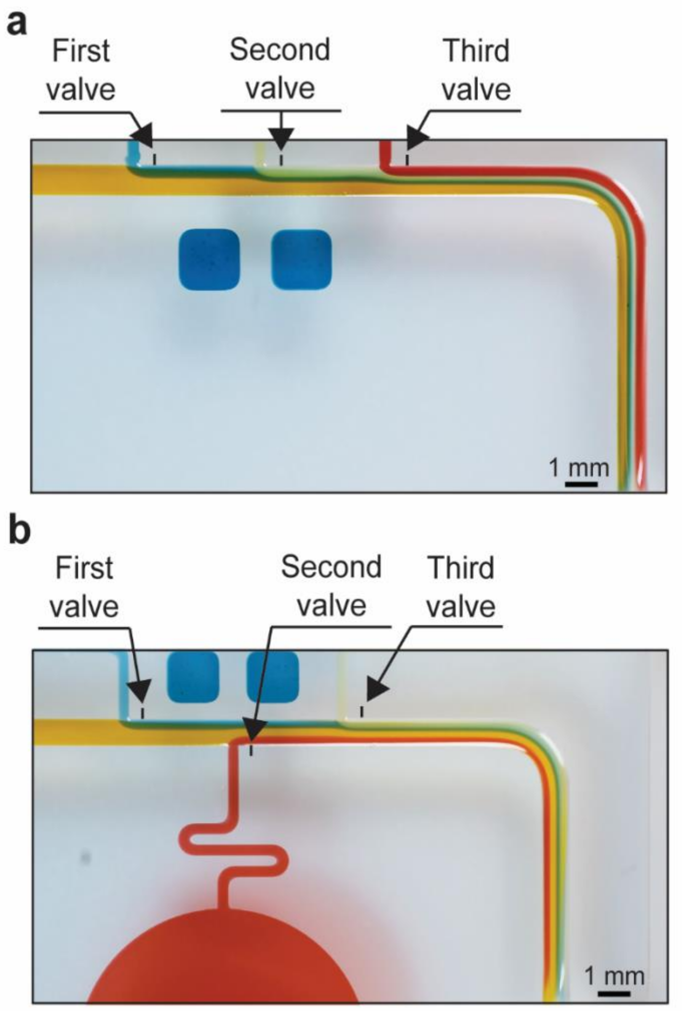

**Figure SI.5.7.** *Geographical location of layer width measurements for each TGV on the same and opposing sides of the main channel.* Layer width measurements were taken 0.5 mm from each TGV channel wall (black lines) for each of the corresponding TGV's colored fluid layer.

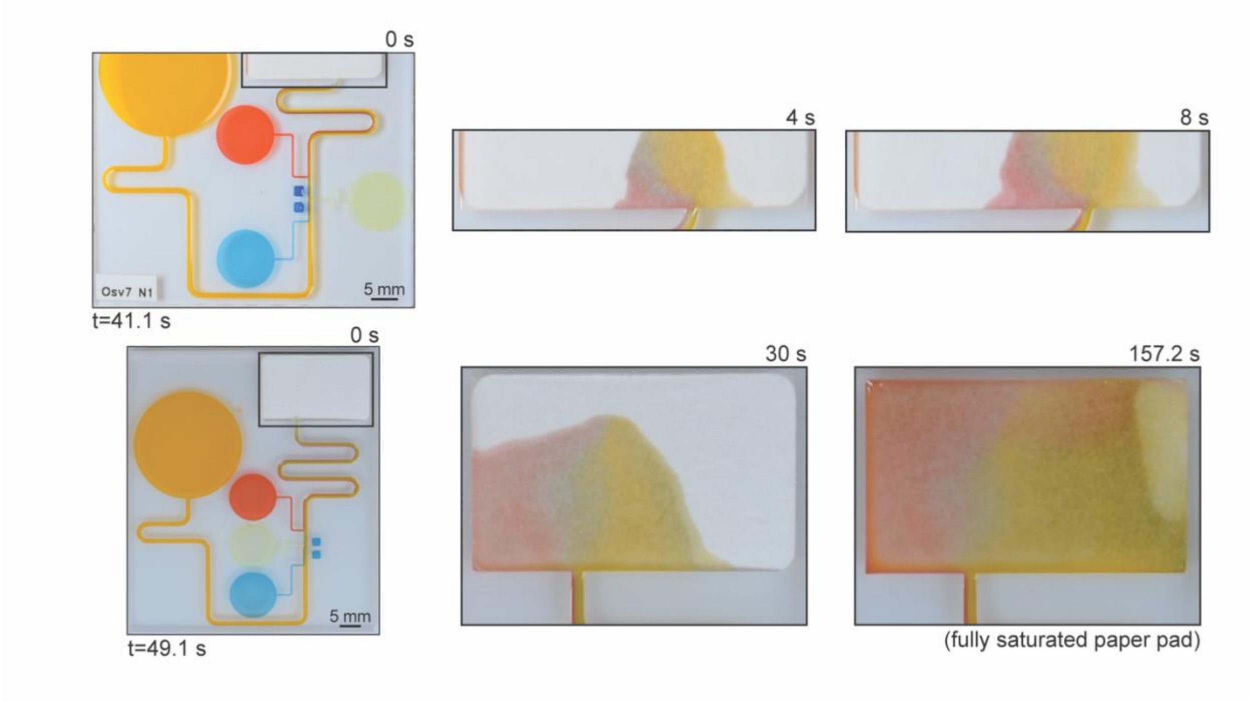

**Figure SI.5.8.** Fluid layers remain partially unmixed after reaching the paper pads. Images of the flow profiles of the paper pads for devices used in Figure SI.5.2 (top) and Figure SI.5.4 (bottom).

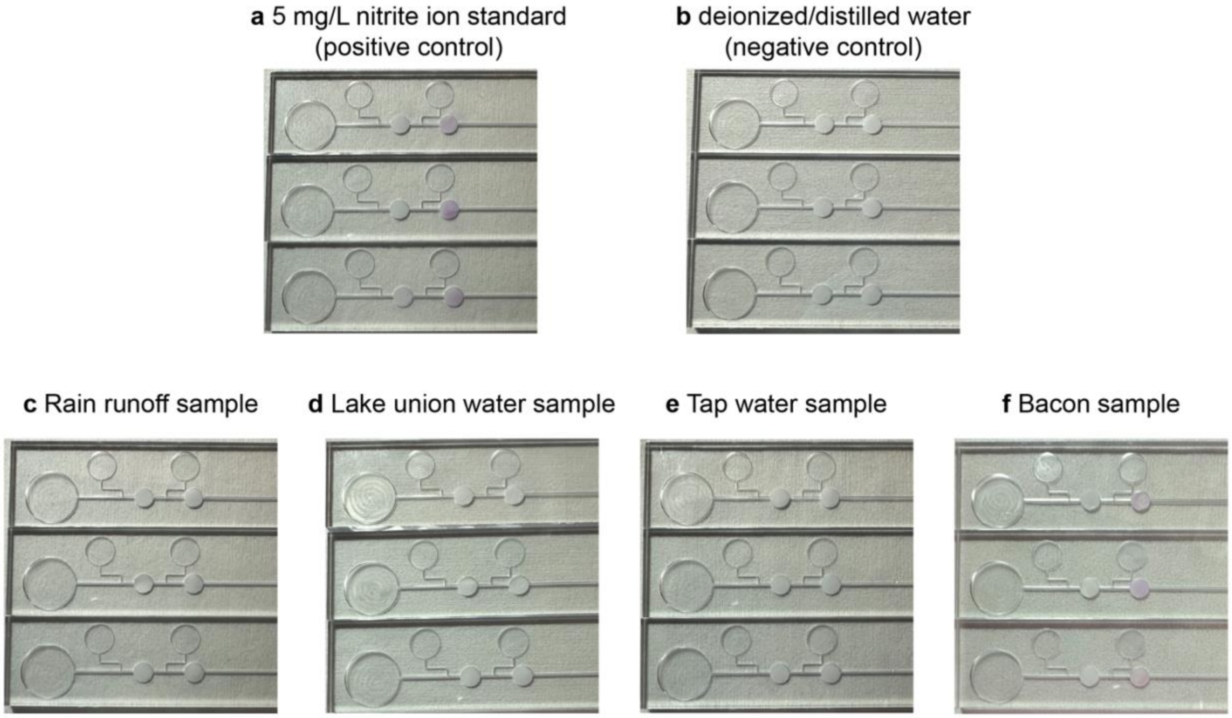

**Figure SI.5.9.** Images of all Griess reaction trials for a 5 mg/L nitrate ion standard (a), deionized/distilled water (b), Lake Union sample (c), rain runoff (d), tap water (e), and a bacon sample (f).

**Table SI.5.1.** Detected nitrite concentrations in environmental and meat samples

| Sample | Mean concentration (± standard deviation) (mg/L) |
|---|---|
| Seattle tap water | ND* |
| Lake Union | 0.1535 (0.3) |
| Rain runoff | ND* |
| Bacon meat | 10.7272 (± 0.5)** |

*Reported as not detected (ND) as the value is below the lowest detectable standard solution.

**Value was obtained by multiplying the diluted bacon sample by 4 (bacon meat sample was diluted at a ratio of 1:4) and is reflected in the table.

#### SI.6. Supporting information references

1. J. Berthier, D. Gosselin, E. Berthier 2015 A generalization of the Lucas–Washburn–Rideal law to composite microchannels of arbitrary cross section *Microfluid Nanofluid* **19**:497-507
2. J.C. Tokihiro, A.M. McManamen, D.N. Phan, S. Thongpang, T.D. Blake, A.B. Theberge, J. Berthier 2024 On the Dynamic Contact Angle of Capillary-Driven Microflows in Open Channels Langmuir **40**, 13, 7215–7224
3. J. Bico, C. Marzolin, D. Quéré 1999 Pearl drops *Europhys. Lett.* **47**(2), 220-226
4. J. Berthier, K.A. Brakke, The Physics of Microdroplets, Scrivener-Wiley Publishing, 2014.
5. D. Quéré 1997 Inertial Capillarity. *Europhys. Lett.* **39** (5), 533−538.
6. J. Berthier, A.B. Theberge, E. Berthier, Open-Channel Microfluidics, Fundamentals and Applications, Second Edition, IOP Publishing, 2024
7. R. Lucas 1918 Ueber das Zeitgesetz des kapillaren Aufstiegs von Flüssigkeiten*, Colloid Polym. Sci*. **23** (1), 15−22.
8. E.W. Washburn 1921 The dynamics of capillary flow. *Phys. Rev.*, **17** (3), 273
9. E.K. Rideal 1922 On the flow of liquids under capillary pressure, *Philos. Mag*. **44** (264), 1152−1159
10. C.H. Bosanquet 1923 On the flow of liquids into capillary tubes. *Philos. Mag. Ser*.**6, 45**, 525−531
11. D. Yang, M. Krasowska, C. Priest, M.N. Popescu, J. Ralston, Dynamics of Capillary-Driven Flow in Open Microchannels *J. Phys.Chem*. C 2011, 115, 18761−18769.
12. F.F. Ouali; G. McHale, H. Javed, C. Trabi, N.J. Shirtcliffe, M.I. Newton, Wetting Considerations in Capillary Rise and Imbibition in Closed Square Tubes and Open Rectangular Cross- Section Channels. Microfluid. Nanofluid. 2013, 15, 309−326.
13. P. Kolliopoulos, K.S. Jochem, D. Johnson, W.J. Suszynski, L.F. Lorraine, S. Kumar 2021 Capillary-flow dynamics in open rectangular microchannels, *Journal of fluid mechanics* **911** A32-1 – A32-23
14. J. Berthier, K.A. Brakke, E. Berthier 2014 A general condition for spontaneous capillary flow in uniform cross-section microchannels *Microfluid Nanofluid* **16**:779–785
15. T.D. Blake, J.M. Haynes 1969 Kinetics of Liquid-Liquid Displacement. *J. Colloid Interface Sci.* **30** (3), 421–423
16. T.D. Blake 2006 The physics of moving wetting lines, J*. Colloid Interface Sci*. **299** (1), 1−13
17. J.C. Tokihiro, Wan-chen Tu, J. Berthier, J.J. Lee, A.M. Dostie, Jian Wei Khor, M. Eakman, A.B. Theberge, E. Berthier 2023 Enhanced capillary pumping using open-channel capillary trees with integrated paper pads *Physics of fluids* **35** (8), p.082120-082120
